# A two-stage algorithm underlies the transformation from vision to familiarity in the primate brain

**DOI:** 10.1101/2025.06.13.659490

**Authors:** Simon Bohn, Catrina M. Hacker, Barnes G. L. Jannuzi, Travis Meyer, Madison L. Hay, Nicole C. Rust

## Abstract

How is the act of seeing an image transformed into the memory that it has been seen? To investigate, we leveraged the systematic variation with which some images are better remembered than others, image memorability, to compare neural responses in inferotemporal cortex (ITC) and the hippocampus (HC) as macaque monkeys performed a single-exposure visual familiarity task. We found evidence for a two-stage algorithmic transformation from visual representations to familiarity, including a previously undescribed computational transformation of familiarity in the medial temporal lobe. At the first stage, more memorable images elicited more vigorous ITC firing-rate responses and stronger ITC familiarity signals (reflected as repetition suppression). This led to a counterintuitive intermingling of familiarity and memorability signals in ITC, but a representation that could be read out by a linear decoder. At the next stage, the medial temporal lobe selectively extracted ITC familiarity signals to produce a more isolated familiarity representation with minimal memorability modulation, reflected downstream in HC. These results shed light on how seeing is transformed into familiarity, and they establish the existence of a previously undescribed medial temporal lobe computation.

## Introduction

Human and non-human primates possess the remarkable ability to quickly, accurately and seemingly effortlessly remember what they have seen before. How the brain supports this behavior is not fully understood (Fig. 1a). One well-established behavioral signature of visual familiarity (also called visual recognition memory) is that some images are consistently more memorable or forgettable than others, a property called image memorability (Fig. 1a-b). Image memorability is consistent across human observers^1–7^ and between humans and rhesus monkeys^8^. Likewise, image memorability is reflected in deep artificial neural networks that are trained to categorize objects with no explicit optimization for memorability^8,9^, suggesting that it is a natural byproduct of a system wired up to see. Here, we used image memorability variation to understand how the brain transforms vision into familiarity.

**Figure 1:**
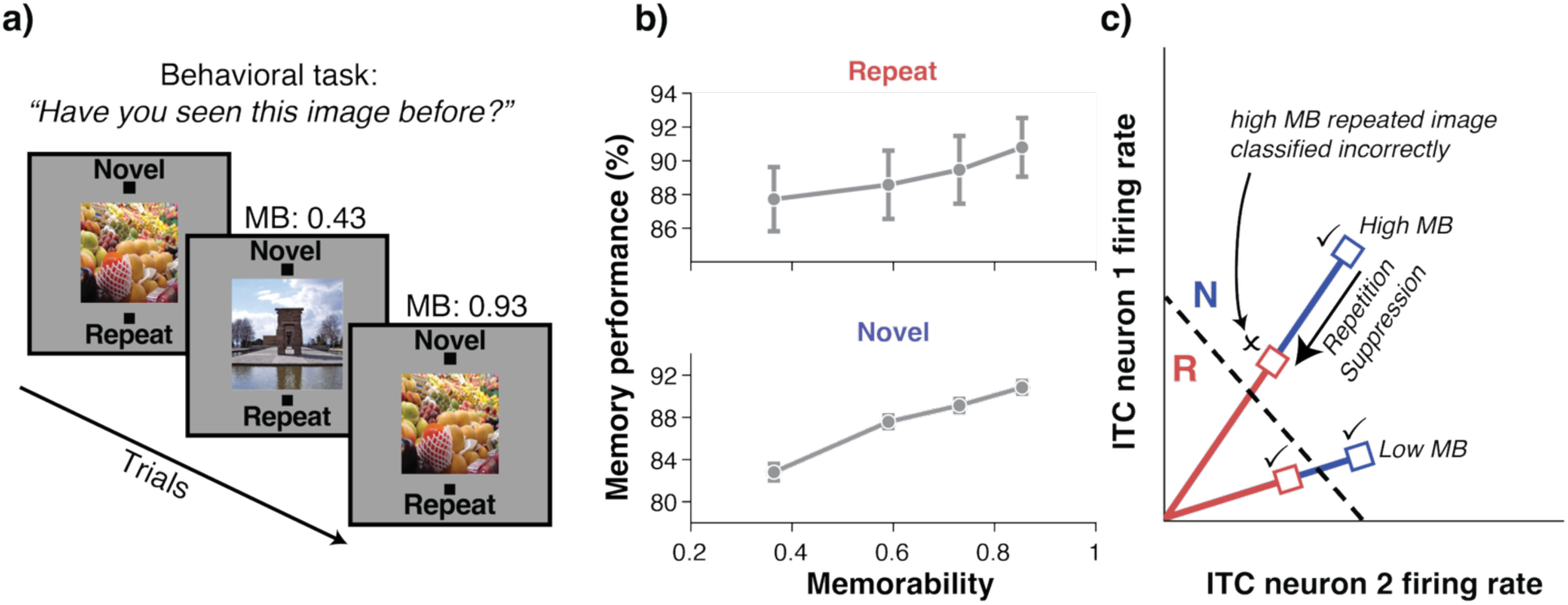
Task, behavior and relationship between familiarity and memorability in ITC. **a)** Summary of the behavioral task. Two rhesus monkeys were shown one image per trial and indicated whether they judged the image to be novel (never seen before) or repeated (seen exactly once before). The number of intervening trials between novel and repeated presentations (n-back) ranged from 1 to 192, with most occurring in the range of 2-64. Images naturally varied in memorability (MB), a scalar (range 0-1) that corresponds to the likelihood of a human correctly remembering that image. **b)** The pooled results of the two macaque monkeys on this task for repeated (top) and novel (bottom) images (n=10,816 trials of each type) binned into four equally sized memorability groups. As predicted from the human-derived memorability scores, higher memorability images were more likely to be correctly recognized as repeated by the monkeys. Error bars depict the standard error of the mean and are larger for repeated behavior compared to novel behavior because “repeat” trials vary in the number of intervening trials since the novel presentation, whereas all “novel” trials are the same. **c**) Schematic of visual familiarity representations in ITC. Axes represent firing rates of two idealized ITC neurons. When viewed, trials induce a unique pattern of firing in ITC (vector direction in this space; each vector represents a different image) represented by the blue vector for the novel presentation. When an image is repeated, the pattern of firing activity produces the same vector angle, but with reduced vigor, so the magnitude of the vector is reduced (red vector), a phenomenon known as repetition suppression. The dashed line shows the decision boundary for a decoder that categorizes trials as novel or repeated based on the overall spiking vigor in ITC, with images evoking population firing greater than the boundary being classified as novel (N) or less than as repeated (R). Checkmarks indicate firing rate patterns that would be correctly classified to correspond with the trial that evoked them while “x” marks those that would not. More memorable images induce higher firing vigor, challenging familiarity decoding schemes based on vigor alone. For example, a decision boundary that correctly classifies low memorability images (dashed line) incorrectly classifies high memorability images, leading to a systematically reversed prediction of behavior as a function of memorability based on a response vigor decoding scheme (Fig. 2b). Characterizing whether and how this contradictory representation in ITC is transformed by downstream brain regions is the subject of this paper.

Several lines of evidence suggest that the signal the brain relies on for visual familiarity is related to repetition suppression, the characteristic reduction in firing to a repeated image, which can be accessed by reading out the overall vigor of a population response^10–16^. For example, a weighted linear decoding scheme based on firing vigor in ITC accurately aligns with subjects’ decaying ability to report familiarity as time passes since seeing an image^15^. However, neural phenomena associated with image memorability challenge whether such a readout might lead to familiarity behavior. Namely, previous work has shown that more memorable images evoke more vigorous responses^8^, meaning that increased firing due to memorability acts in opposition to the decreased firing due to familiarity. This leads to the challenge illustrated in Fig. 1c where a firing threshold that does a good job differentiating novel from familiar images for a low memorability image does a poor job with a high memorability image. This results in systematically reversed predictions of behavior as a function of memorability when using firing vigor to decode familiarity status (Fig. 2b).

**Figure 2:**
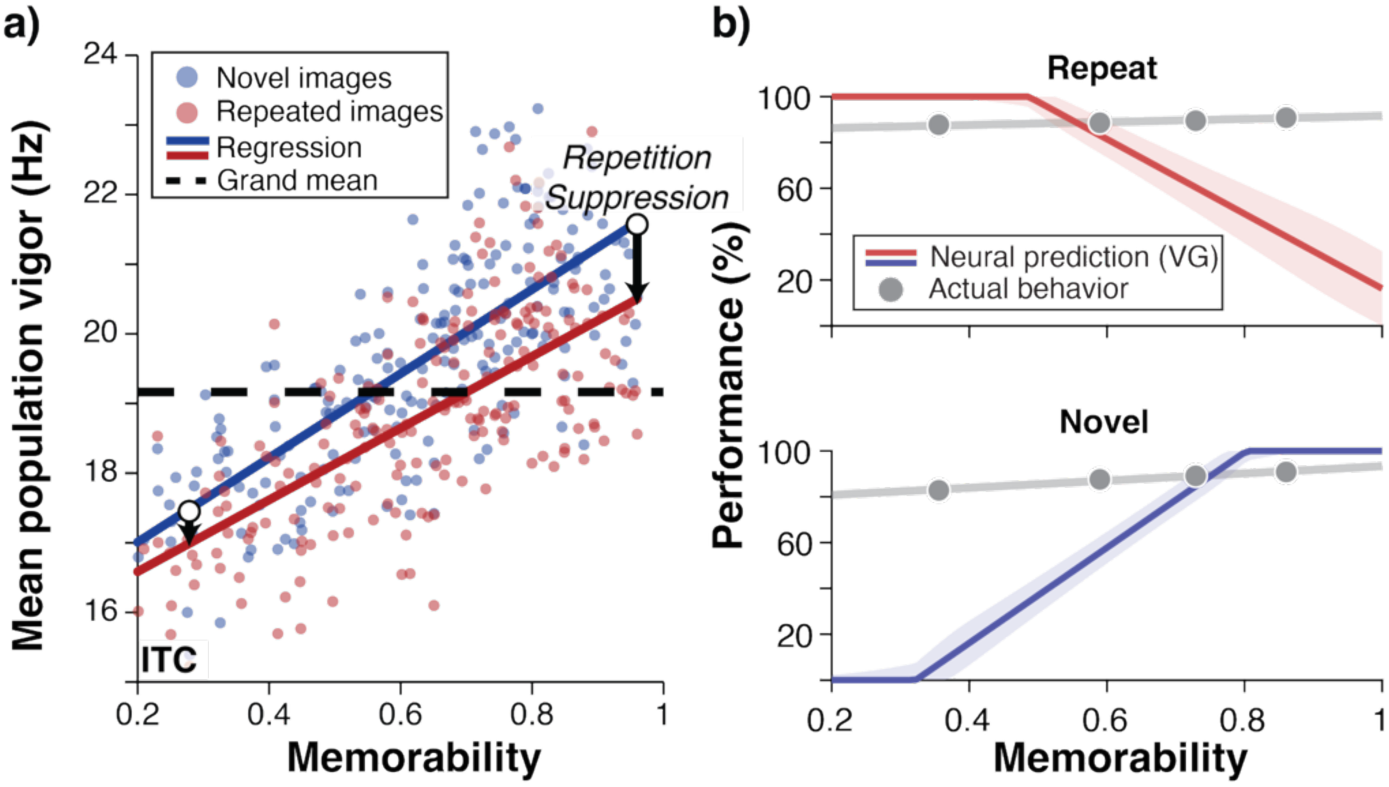
Higher memorability leads to greater repetition suppression in ITC, but also a counterintuitive representation in which it is not clear how familiarity is decoded to produce the observed familiarity behavior. **a)** Neural data recorded in ITC with spikes counted in the 100-500ms window following image presentation. Each dot represents mean population vigor, the firing rate averaged across all units (n=623 units) to a novel (blue) or repeated (red) image of a given memorability. Grand mean firing rate across all units and images is shown as a dashed line; this line is the decision boundary in the linear decoder shown in panel (b). The solid lines are linear regressions of memorability versus firing vigor for novel and repeated images, respectively. Black arrows indicate the difference between novel and repeated firing (repetition suppression, RS), which increased with memorability (Fig. S3). **b)** Cross-validated linear familiarity decoder based on reading out the population vigor applied to the data in panel (a), weighting each unit positively and equally. In this vigor (VG) decoder, a single decoding parameter (the decision threshold) was trained on a subset of images and then tested on different subsets of images binned by memorability. Note the mismatched neural prediction (red, repeated images, and blue, novel images) from the actual behavior (gray). Neural predictions are rescaled with a multiplicative factor, consistent with an adjustment in population size (see Methods). Error shadows depict one standard error.

Therefore, despite previous reports^10–16^, it is not clear whether and how the brain uses ITC visual representations to make familiarity judgments. While it may be tempting to propose that the brain accesses repetition suppression by itself (the firing rate difference between novel and repeated images, compared to raw vigor described above) to drive familiarity judgments, such explanations are circular, as they require familiarity to explain familiarity (“What was the firing rate of *this* image the last time *it* was seen?”). As such, this raises the question of how the brain differentiates between firing rate modulations due to familiarity and memorability.

An analogous conundrum has been described for low-level visual features like image contrast, where the signs of visual and familiarity modulations also operate in opposition (that is, contrast increases firing but familiarity decreases it). One study reconciled this by demonstrating that familiarity signals could be selectively extracted from ITC while discarding fluctuations attributed to image contrast with a linear decoder specialized for this purpose to accurately predict contrast-invariant familiarity behavior^17^. However, it’s unclear whether the proposed contrast mechanism applies to memorability, given that familiarity performance is unaffected by image contrast (familiarity reports are similar for low-versus-high contrast images that elicit lower-versus-higher ITC vigor), yet familiarity performance consistently increases with image memorability. Furthermore, that study proposed how ITC could in principle be read out, but it did not provide empirical evidence that such a process indeed takes place (e.g. from measurements downstream of ITC in the medial temporal lobe, where repetition suppression has also been observed^10,13,18–22)^.

In this study we targeted two questions: First, can isolated familiarity signals be extracted from ITC despite the intermingling of familiarity and memorability that follows from opposite sign modulations? Second, is there evidence of such an extraction taking place? We found evidence of a two-stage algorithmic transformation from vision to familiarity within ITC and the medial temporal lobe, the results of which are reflected in the hippocampus (HC). In the first stage, higher memorability images generated stronger firing rates in ITC, and larger familiarity signals for more memorable images. While this produced a counterintuitive representation of familiarity in ITC, the mapping from the ITC population response to behavior could be accounted for by a specialized linear decoding scheme. In the second stage, a computational transformation in the medial temporal lobe largely removed memorability modulation, consistent with the proposed ITC linear decoding scheme. These findings refine our understanding of how memorability and familiarity are multiplexed in ITC and suggest the existence of a previously undescribed computation in the medial temporal lobe that transforms visual familiarity information arriving from ITC.

## Results

### Single-exposure visual familiarity task

Two rhesus macaque monkeys performed a single-exposure visual familiarity task^15^ in which they sequentially viewed images varying in memorability and reported after each trial whether the image was novel (never seen before in this or any previous sessions) or repeated (seen exactly once before in this session; Fig. 1a). They were rewarded with juice for correct reports. Monkeys initiated a trial by fixating on a central fixation point. After fixating for 500ms, the monkeys maintained fixation while an image appeared for 500ms, subtending 4 degrees of the central visual field. After the image disappeared, the monkeys indicated whether the image was novel or repeated by making a saccade to one of two response targets. Correct responses were rewarded with juice. All images were shown exactly twice, and the time between the novel and the repeated presentation of an image varied by the number of intervening trials (n-back) from seconds (one intervening trial) to minutes (192 intervening trials; see Methods).

Images depicted a diverse range of object categories and scenes and spanned a broad range of memorability scores (Fig. S1). Each image was assigned a memorability score ranging from 0 to 1 reflecting how likely an image is to be remembered^7^ as quantified by a model trained to predict human familiarity behavior with accuracy near the ceiling imposed by inter-subject consistency (MemNet^5^). Consistent with previous reports^8^, human-based memorability scores predicted familiarity performance for monkeys: In the data pooled between both monkeys a logistic regression showed a significant relationship between memorability and performance (for repeated images, *β*=0.80, odds ratio = 2.24, p = 5.4e-8; for novel images, *β* =1.5, odds ratio = 4.57, p = 3.96e-28; n = 10,816 trials for each type, Fig. 1b). Memorability affected behavior in the same way when each monkey’s data was analyzed individually (Fig. S2).

### Neural recordings

As the monkeys performed this task, we recorded neural activity in ITC or HC with a 24-channel recording probe lowered acutely for each session. We included units with activity that reflected task-responsiveness (i.e. changed their firing rate during the trial compared to baseline; M1: n=290 ITC and 312 HC units; M2: n = 333 ITC and 503 HC units). Because the responses of individual ITC and HC units are often tuned for specific visual features, estimating the overall population response requires many hundreds of units. Therefore, we concatenated units across sessions into a pseudopopulation and aligned images of similar memorability into “pseudoimages” as in refs^8,15–17^. All neural analyses were performed on these pseudoimages, to which we refer interchangeably with “images” when describing analyses of these data. When creating the pseudopopulations, we aligned trials in a manner that preserved familiarity, n-back, and approximate memorability (see Methods).

### Memorability and repetition suppression are correlated in ITC

While memorability is known to influence ITC population firing vigor^8^, how it systematically relates to the magnitude of repetition suppression has not been previously documented. We thus characterized the firing vigor (the mean firing rate of the neural population to an image across all units) to novel and repeated presentations of an image as a function of memorability. Images with higher memorability scores were associated with more repetition suppression (Figs. 2a and S3). This can be seen in Fig. 2a, where the amount of repetition suppression is visualized with black arrows between the novel (blue) and repeated (red) lines. Images with higher memorability (the longer right arrow) evoked more repetition suppression than images with lower memorability (the shorter left arrow). Likewise, the repetition suppression of individual images, quantified as the difference in firing rate between the population response to their novel and repeated presentation, was significantly correlated with its memorability (Fig. S3, r(206) = 0.23, p = 0.001). This was also the case when each monkey’s neural population was analyzed individually (Figs. S3-S4). These results establish, for the first time, that memorability is associated with the strength of a putative familiarity signal in ITC.

### Memorability interferes with vigor-based decoding of familiarity

To understand the interaction between familiarity and memorability signals in ITC, we quantified their relative modulations of overall firing rate. Consistent with previous reports^8^, memorability was positively correlated with ITC population firing vigor. We found that this was true for both repeated and novel images (Fig. 2a: repeated images Pearson’s r(206) = 0.70, p=6.7e-32, novel images r(206) = 0.74, p=4.9e-38. See Fig. S4 for these data analyzed in each monkey individually with similar results.) These strong correlations translated to a 19.2% difference in firing vigor between novel images of low (10^th^ percentile) versus high (90^th^ percentile) memorability. In comparison, the modulation of firing by familiarity translated to a 4.0% difference between firing vigor to novel versus repeated images (across all memorability scores). In other words, memorability modulation was ∼5-fold larger than familiarity modulation, dominating population vigor.

The intuition that follows from Fig. 2a is that memorability modulation will interfere with attempts to decode familiarity from vigor in ITC (i.e., as illustrated in Fig. 1c). To test if that intuition held in these data, we trained a cross-validated total spike count linear decoder, which we called the vigor (VG) decoder, to distinguish novel from repeated images on the basis of their response magnitude. Specifically, the VG decoder weighted each unit equally and positively to classify the neural responses to each image as novel or repeated based on whether the firing rate fell above or below the decision boundary (dashed line in Fig. 1c and 2a).

When applied to the neural data we observed in ITC, this decoder produced predictions for repeated images that were strikingly different from observed behavior (Fig. 2b). An alternative linear decoding scheme optimized for familiarity performance that allowed for variable weighting of each unit (and negative weights to account for units that might exhibit increased firing to repeated images) produced qualitatively similar misaligned predictions (Fig. S5). These results follow from the intuition sketched in Fig. 1c that, despite its success in predicting other familiarity behaviors, previous methods of linearly decoding ITC cannot account for memorability behavior. We therefore turn to the question of whether a different linear decoding scheme applied to ITC can produce more accurate behavioral predictions.

### Stage 1: A linear decoding scheme can selectively extract familiarity signals from ITC to predict memorability behavior

Because reading out ITC firing vigor was insufficient to explain memorability behavior, we investigated whether memorability and familiarity were reflected in the ITC population in a way that could support behavior via a different linear decoding scheme. The fact that modulations due to familiarity and another image-related factor that modulates population vigor, contrast, can be disambiguated with a linear decoding scheme^17^ suggests the (as yet untested) hypothesis that a similar scheme could work for memorability. This is not a given because memorability is a different visual feature and differs from contrast in that familiarity behavior is sensitive to memorability, but it is contrast-invariant. In this and the next section, we test predictions of and find support for the two-part hypothesis that (1) a representation where memorability behavior is linearly extractable from ITC exists, and (2), the result of such an extraction is reflected downstream of ITC, in HC.

The hypothesized linear decoding scheme relies on representations of familiarity and memorability in ITC’s firing being at least partially nonoverlapping, allowing familiarity to be extracted partially independently of memorability. It therefore cannot work if they completely overlap, such as if memorability and familiarity exerted the same influence on the activity of each neural unit. At the other extreme, linearly decodable information about memorability and familiarity could be completely non-overlapping (i.e., orthogonal), allowing one to be extracted independent of the other. These two scenarios form the ends of a continuum that spans from complete through partial overlap to none. In question are whether memorability and familiarity are sufficiently non-overlapping to extract familiarity independent of memorability with minimal loss of familiarity information, and whether this can be done in a way that aligns with observed memorability behavior.

Consistent with memorability and familiarity being partially separable, a linear decoder axis for memorability (MB, which separated high and low memorability images) and a decoder axis for total population vigor (VG, the same population vigor decoder used in Fig. 2b that weighted each unit positively and equally) were partially overlapping in ITC (mean +/− se: 51.6 +/− 1.4 deg, M1: 49.7 +/− 2.2 deg, M2: 51.6 +/− 2.0 deg, Fig. 3a and S6-S7). Because the axes were not completely overlapping, this suggested that some information about familiarity should be extractable independent of memorability, but because the axes were not orthogonal, some information would be lost. This raises questions about whether the ITC representation can be linearly decoded to align with memorability behavior, and, if so, how much familiarity information is preserved by that decoder.

**Figure 3:**
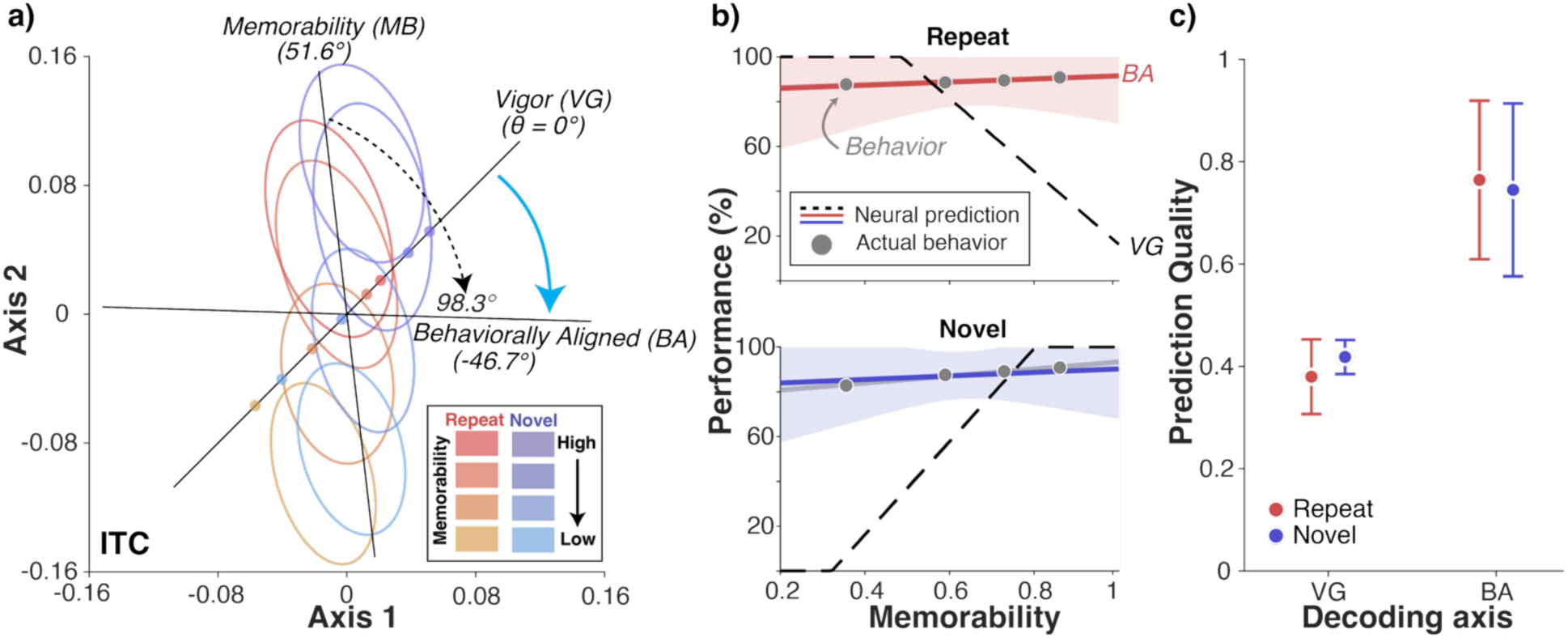
Memorability behavior is linearly decodable from ITC. **a)** Projections of ITC neural data onto a plane defined by an axis corresponding to the repetition-suppression-based vigor decoder (where every unit is weighted equally, ‘Vigor (VG)’) and a linear memorability decoding axis optimized to decode neural firing to high versus low memorability images, ‘Memorability (MB)’. These two axes fell 51.6 degrees apart, establishing that memorability is partially overlapping with overall spiking vigor in ITC (where nonoverlapping is indicated by orthogonality, 90 degrees). Ellipses indicate the one-standard-deviation contour of the projections of ITC population responses onto this plane. The eight ellipses correspond to novel (blues) and repeated (reds) images grouped by memorability scores into quartile bins (hue). Translucent dots are projections of the means of each memorability/familiarity bin onto the vigor axis and give an intuition of how these axes are read out as linear discriminators. Here, the data are centered such that the decision boundary is at the origin, so that positive values decode as novel and negative as repeated. The most memorable repeated images (dark red) falling on the “novel” side of the decision boundary are consequently classified incorrectly, corresponding to the poor prediction quality of the VG decoder (Fig. 2b and Fig. 3b-c). Rotated axes in this plane reflect linear combinations of the VG and MB axes. To test if familiarity could be extracted from population vigor despite memorability modulation, we swept through possible angles from the VG axis and found the decoding axis on this plane that best maps to behavior (‘Behaviorally aligned (BA)’), which fell just beyond orthogonal to the MB axis, at 98.3 degrees clockwise (dashed black arrow), or equivalently 46.7 degrees clockwise of the VG axis (cyan arrow). Marked angles reflect means across 1000 cross-validation iterations. **b)** Comparison between observed behavior (gray) and neural predictions for the vigor decoder axis (VG, dashed line; copied from Fig. 2b) and behaviorally aligned decoding axis (BA, red and blue lines correspond to repeated and novel behavior, respectively). Data used to make these predictions were held out of the sets used to train the memorability decoder and optimize the angle of the BA decoder axis, and the solid line shows the average predicted behavior across 1000 cross-validation iterations. Error shadows depict the standard error across cross-validation iterations. Neural predictions are rescaled with a multiplicative factor consistent with an adjustment in population size (see Methods). **c)** Comparison between prediction quality when decoding behavior using the vigor (VG) axis versus the behaviorally aligned (BA) axis using the same fully held-out data as described above. Error bars depict standard error. The spike counting window throughout this figure was the same as Fig. 2 (100-500ms).

To test if memorability’s influence on the population vigor could be isolated and selectively removed so that familiarity predictions aligned with behavior could be decoded, we examined decoders on the plane formed by the linear decoding axes for memorability and population vigor described above. For a visualization of how this plane is derived, see Fig. S8. Weighted linear combinations of the MB and VG axes constitute a new linear decoder axis which are instantiated by sweeping through rotations within this plane. The axis orthogonal to MB (90 degrees away) in this plane is special, as it reflects the extraction of familiarity information independent of memorability.

Any axis in this plane can be evaluated for its ability to map to observed memorability behavior by using it to decode whether the neural firing evoked by each image corresponded to a novel or repeated presentation, then comparing those predictions with actual behavior. These predictions were quantified with a measure of prediction quality (PQ, see Methods) to evaluate the quality of a decoder’s prediction with a scale that ranged from 0 (orthogonal to the behavioral slope) to 1 (exactly matching). Applying this measure of prediction quality to the vigor decoder (initially shown in Fig. 2b and copied onto Fig. 3b as a dashed line) quantifies the poor alignment that comes from attempting to use vigor alone to decode memorability behavior (mean PQ +/− se for repeated behavior: 0.38 +/− 0.07; for novel behavior 0.42 +/−0.03, and similar results for each monkey analyzed individually, Figs. S6-S7).

We examined the family of linear decoder axes that populated the plane formed by the MB and VG axes by sweeping through all possible rotations in incremental steps. Our approach uses one set of training data to find the memorability axis, a second set of training data to find the best behaviorally aligned familiarity-decoding angle on the MB/VG plane, and fully held-out test data to measure behavioral performance predictions along this decoding axis (see Methods and Fig. S8). In other words, our approach applies a two-step procedure to identify the linear decoding axis using training data and a third step to evaluate performance along that axis with fully held out test data.

We found that an axis that lay just beyond orthogonal (mean +/− se 98.3 +/− 5.6 deg, M1: 99.0 +/− 10.3 deg, M2: 97.4 +/− 7.5 deg) to the memorability decoder axis best aligned with behavior as quantified by having the maximum prediction quality. We termed this axis the behaviorally aligned (BA) axis. Intuitively, the angle it landed at makes sense because 90 degrees would correspond to familiarity modulations with minimal stimulus-evoked memorability influence on the firing vigor, so the best possible prediction quality should fall just past orthogonal to align with higher familiarity performance for more memorable images.

Decoding along the BA axis produced good predictions of both repeated and novel behavior when tested on fully held-out data that was not used to train either the memorability decoder or to find the BA axis (repeated-behavior PQ mean +/− se: 0.76 +/− 0.15, novel: 0.75 +/− 0.17, Fig. 3c.) This finding was replicated when each monkey was analyzed individually (Figs. S6-S7). A bootstrap analysis (see Methods) revealed that the BA decoder significantly outperformed the VG decoder in predicting memorability behavior (ΔPQ and 95% CI for repeated behavior: 0.39, [0.36, 0.42], H0: ΔPQ=0, p < 1e-4, for novel behavior: 0.31, [0.28, 0.34], H0: ΔPQ=0, p < 1e-4 and similar results when each monkey was analyzed individually, Fig. S6-S7).

Although our procedure uses cross-validation to prevent overfitting, identifying the behaviorally aligned decoder axis relies on fitting to behavioral data, thus raising an important question: are these results inevitable? Or, in other words, what is the null model? Here, the null model is a population that lacks a population geometry that allows for reliable identification of the BA axis. By extension, the most insightful measure for comparison is the precision of the BA decoder axis angle relative to the vigor axis across cross-validation splits. To create such a null model for comparison, we shuffled memorability scores by randomly reassigning them to novel/repeated image response pairs on each cross-validation iteration before training and testing. In the intact data, decoding angles were highly consistent: in the pooled population the mean +/− se was –46.7 +/− 4.9 deg, in M1: −49.3 +/− 10.0 deg and M2: −45.7 +/− 6.7 deg. In contrast, null model decoding angles varied wildly, creating a near uniform distribution across all possibilities from −90 to 90 degrees (mean +/− se 3.2 deg +/− 44.5 deg, Fig. S9). These results confirm that the identification of the BA axis is not an inevitability of our procedure for identifying it but a product of the structure of the neural data.

These results establish that the ITC population geometry supports the ability to linearly decode memorability behavior, in principle. As a final consideration, we considered whether sufficient information about familiarity survives this decoding transformation to make this a plausible account in practice. For instance, if this scheme destroyed 95% of all familiarity information, it would be a dubious proposal. To evaluate this issue, we compared total familiarity performance for the behaviorally aligned linear decoding axis with other decoders on the plane (Fig. S6-S7, S9). Compared to decoding using population vigor alone, decoding along the BA axis retained substantial amounts of (or increased) linearly available familiarity information: in the pooled data performance was 1% higher than decoding at the VG axis, in M1 62% of performance was maintained and in M2, performance increased by 35%.

Altogether, these results suggest that a linear readout along the behaviorally aligned axis can plausibly explain how reflections of familiarity and memorability are configured in ITC to support behavior. Specifically, although familiarity and memorability influence population vigor in an overlapping manner, memorability behavior can be linearly decoded from ITC via a decoding axis just past orthogonal to memorability modulation with minimal loss of familiarity information. This raises the next question: is there any evidence that the brain performs this algorithmic transformation?

### Stage 2: The hippocampus reflects medial temporal lobe computations that extract ITC familiarity signals

Our hypothesis for the second stage of this transformation is that familiarity information will be reflected downstream in a manner consistent with the behaviorally aligned decoding scheme described above, and makes three specific predictions. First, the format of the familiarity signal should remain as repetition suppression. This prediction follows from the fact that we measured the angle between the vigor and behaviorally aligned axis as acute (−46.7 deg, Fig. 3a), coupled with the fact that repetition suppression should remain along any decoding axis with an acute angle relative to the vigor axis (i.e. one that does not invert the direction, see Fig. S10 for a visualization). Second, population firing vigor downstream should have reduced modulation by memorability relative to ITC given that eliminating memorability modulation is the goal of the computation (or equivalently, the behaviorally aligned axis lies nearly orthogonal to memorability in ITC). Third, because the previous two predictions create a population geometry in HC with lessened memorability modulation, reading out HC with the same logic as we applied to ITC should yield a behaviorally aligned axis in HC with a smaller angle relative to the vigor axis as compared to ITC.

To test whether these predictions were realized via computation within the medial temporal lobe (MTL), we recorded from the endpoint of the MTL, the hippocampus (HC), where we found evidence in support of all three predictions. We emphasize that our approach is designed to capture the sum total of all computation that occurs along all stages of the MTL through which signals travel after leaving ITC before their reflection in HC, including perirhinal cortex, parahippocampal cortex, entorhinal cortex and the hippocampus itself. Our approach thus tests whether this computation happens in the MTL but does not pinpoint exactly where.

To test the first prediction of the behaviorally aligned decoding scheme, that repetition suppression to familiar images would be present in HC, we began by comparing the peri-stimulus neural response to images in HC compared to ITC. This revealed that the hippocampus, on average, also exhibited repetition suppression (Fig. 4a and S12). To understand the hippocampal representation in more detail, we computed signed sensitivity to familiarity for each unit (d’, which can be thought of as a normalized measure of how repetition of an image affected a given unit’s firing vigor) and compared the distributions of units in ITC and HC (Fig. 4b). Histograms of d’ in both brain areas were roughly normal, but shifted towards positive values, indicating that both populations were dominated by repetition suppression (For ITC, mean = 0.105, s.d. = 0.11 which differed significantly from 0, t(622) = 23.46, p = 1.1e-87; for HC mean = 0.063, s.d. = 0.12 which differed significantly from 0, t(814) = 14.65, p = 2.2e-43). This was also the case when each monkey’s data was analyzed individually (Fig. S11). To determine if the putative repetition-enhanced units in Fig. 4b were contributing to familiarity information in HC as opposed to appearing enhanced due to noisy responses, we performed a ranked decoder analysis (see Methods) and found that in both ITC and HC, only repetition-suppressed units carried reliable familiarity information (Fig. S11). These results support the prediction (following from the geometry of the readout in ITC described above) that the format of familiarity in hippocampus is repetition suppression.

**Figure 4:**
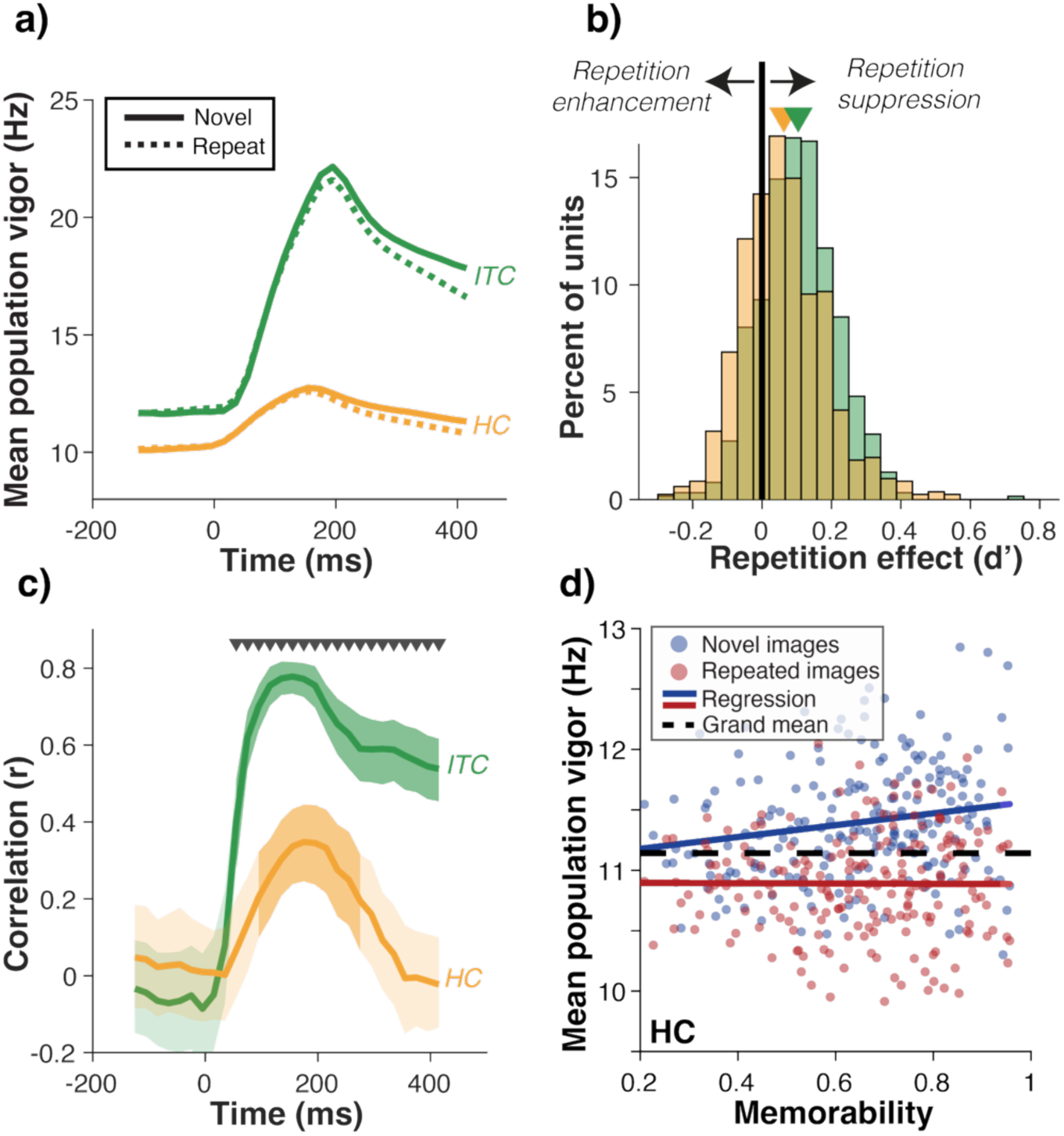
Testing predictions that the medial temporal lobe extracts familiarity information from ITC using the scheme in Fig. 3. **a)** Peri-stimulus neural response aligned to stimulus onset in ITC (green) and HC (orange). Solid lines show the population vigor (firing rate) to novel images and the dashed line to repeated. Data are plotted in a 150ms moving window with 20ms steps. **b)** The distribution of unitwise repetition effect (sensitivity to familiarity, measured in d’) across units in the ITC and HC populations (623 units in ITC, 815 in HC). On the x-axis, a positive value for repetition suppression corresponds to units that fire less to a repeated presentation than to a novel one. The solid black vertical line separates units that were enhanced by repetition (left) from those that were suppressed (right). Triangles point to the mean of each distribution. Familiarity took the form of repetition suppression in HC, as in ITC, supporting the first prediction of the behaviorally aligned decoding scheme. **c)** Pearson’s correlation between memorability and mean population vigor to repeated images as a function of time (150ms moving window with 20ms steps). In ITC (green), this correlation increased rapidly and remained high throughout the viewing window. In HC, this correlation increased transiently followed by attenuation. Error shadows depict 95% confidence intervals bootstrapped across the units. Darker shaded areas were significantly different from no correlation (p < 0.05, corrected for multiple comparisons across time bins) and the gray triangles indicate timepoints where ITC and HC were significantly different (p < 0.05, corrected for multiple comparisons across time bins). **d)** Neural data in the 300-500ms window recorded in HC while monkeys performed the visual familiarity task. Each dot represents the spiking vigor of the population across all units (n=815 units) to a novel (blue) or repeated (red) image of a given memorability. Average firing rate across all units and images is shown as a dashed line; this line would be the decision boundary in the decoder based on population vigor such as the “vigor” decoder described in Fig. 1c. Memorability is attenuated in HC compared ITC (compare with Fig. 2a), supporting the second prediction of the behaviorally aligned decoding scheme.

To test the second prediction of the behaviorally aligned decoding scheme, that memorability modulation should be reduced downstream of ITC, we examined the relationship between memorability and HC spiking vigor for each image as a function of time after stimulus presentation. Unlike ITC, which reflected an immediate and strong correlation throughout the viewing window, in HC, this correlation was attenuated, and by the end of the window, essentially absent (Figs. 4c and S12). By the 300-500ms window (shown in detail in Fig. 4d), the correlation between memorability and firing vigor for repeated images in HC was statistically indistinguishable from no relationship (in the pooled data, r(206) = −0.01, p = 0.935; in M1 r(206) = −0.14, p = 0.04; in M2 r(206) = 0.11, p = 0.13). Likewise, it was not distinguishable from no relationship or modestly positive for novel images (in the pooled data r(206) = 0.20, p = 0.004; in M1 r(206) = −0.05, p = 0.48; in M2 r(206) = 0.28, p = 3.43e-5).

Comparing ITC with HC using a permutation test, memorability modulation of spiking vigor was significantly attenuated in HC as compared to ITC quantified as a reduction in slope for both repeated (ITC β = 5.15, HC β = −0.01, H0: Δβ = 0, p = 5.1e-3) and novel (ITC β = 6.05 and HC β = 0.49, H0: Δβ = 0, p = 4.2e-3) images. Because ITC and HC populations are matched for total familiarity information (see Methods), their differences in memorability cannot simply result from reduced signal or data quality in HC compared to ITC. The significant attenuation of memorability modulation in HC relative to ITC was also present in each monkey individually (Figs. S12-13). Altogether, this reduction in the modulation of spiking vigor by memorability in HC was consistent with predictions of the behaviorally aligned decoding scheme, resulting from a readout of ITC along the behaviorally aligned axis identified in Fig. 3.

To test the third prediction of the behaviorally aligned decoding scheme, that the axis that best reads out behavior in HC is closer to the vigor axis in HC as compared to ITC, we applied the same linear decoding approach taken for ITC (Fig. 3) to HC. Unlike ITC, where memorability largely overlapped with vigor (an approximately 51 degree angle), memorability was nearly orthogonal to vigor in HC (Fig. 5a, mean +/− se 85.2 +/−1.8 degrees, in M1 95.3 +/− 3.0 deg and M2 80.2 +/− 2.5 deg). Likewise, consistent with the third prediction, the behaviorally aligned axis (Fig. 5b) was closer to the vigor axis in HC (in the pooled data, mean +/− se −13.0 +/− 23.0 deg, in M1: 8.6 +/− 37.0 deg, in M2: −19.9 +/− 27.7 deg as compared to ITC at approximately −45 deg). Like ITC, readout along this axis came at minimal cost to familiarity information (Fig. S14).

**Figure 5:**
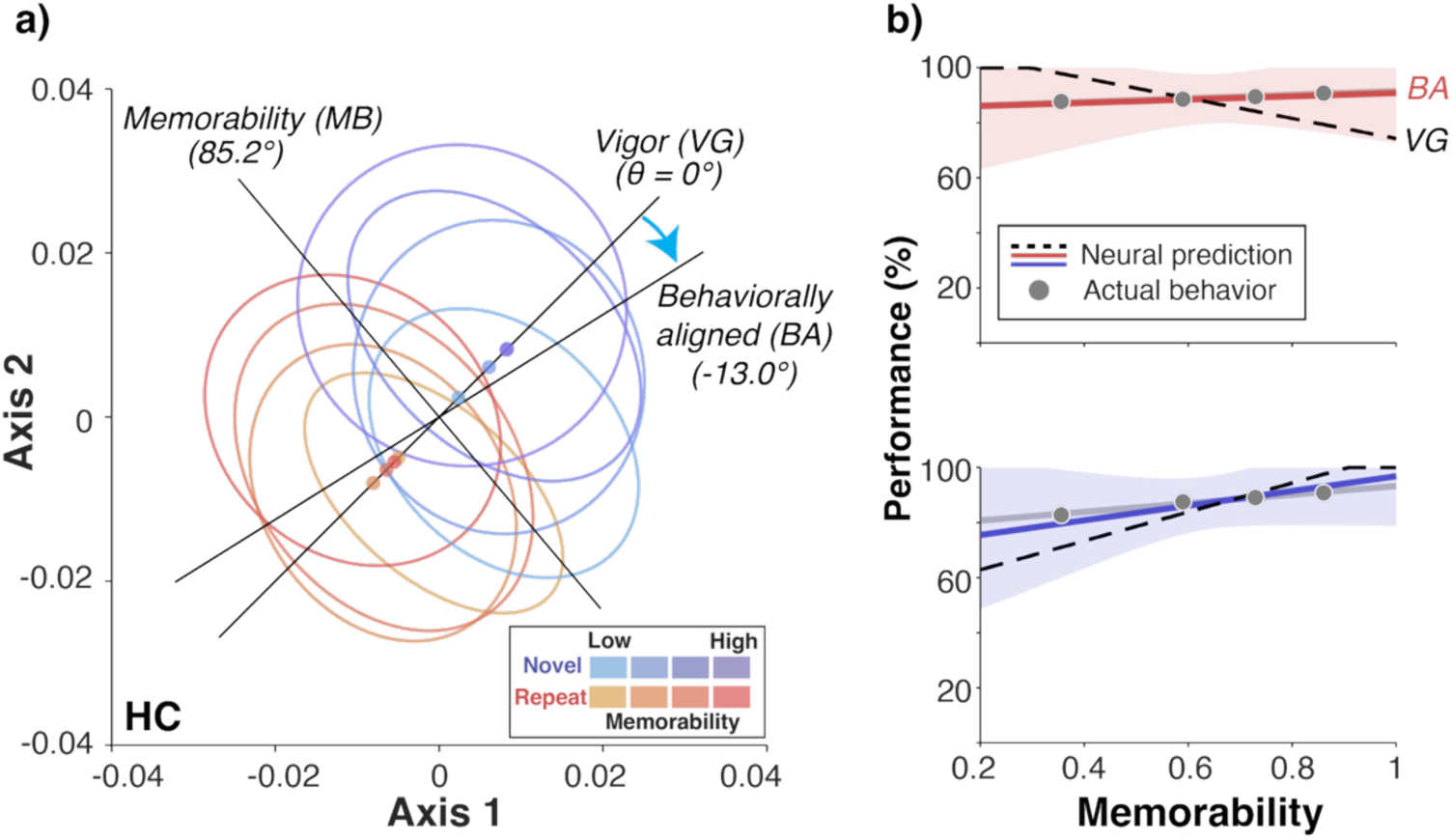
HC population geometry reflects a readout of familiarity along the ITC behaviorally aligned axis. **a)** Projections of HC neural data onto a plane defined by the vigor decoder (where every unit is weighted equally) and the memorability decoder (optimized to decode high versus low memorability images) plotted with the same conventions as Fig. 3a. Ellipses indicate the one-standard-deviation contour for 2D histograms of the projections of HC population responses onto this plane. The eight ellipses correspond to novel (blues) and repeated (reds) images grouped by memorability scores into quartiles (hue). Translucent dots are projections of the means of each memorability bin onto the vigor decoder axis; that the red and blue dots are now cleanly separated along the vigor axis (compared to when they were mixed in ITC) reflects the transformation between the two areas. Also shown is the decoder on this plane with the best ability to map neural data to behavior (behaviorally aligned axis). In contrast to ITC (Fig. 3a), where the MB and vigor axes were overlapping, in HC they are nearly orthogonal, and the behaviorally aligned axis falls closer to the vigor axis (cyan arrow). **b)** Comparison of behavior (gray) with neural predictions at the behaviorally aligned decoding axis (BA, solid red for repeat, blue for novel) and at the vigor (VG, dashed black). Neural predictions are rescaled to the same range as behavior with a multiplicative factor, consistent with an adjustment in population size (see Methods). Across this figure, spikes were counted in the 300-500ms window. Predictions shown in b are the mean of fully held-out sets (i.e., not used to train either the memorability decoder or to determine the location of the behaviorally aligned axis) across 1000 cross-validation iterations.

Together, this supports a picture in which the HC representation is consistent with a readout of ITC along an axis close to its behaviorally aligned axis, and consequently less correction for memorability is required to read out behavior from HC than ITC.

Stepping back, the model tested here focused on transformations of neural representations between brain areas and holds explanatory power at the level of algorithm^23^. The specific representations we characterized were recorded in ITC and HC, but the representational challenge and solution is more general. To make this case, we begin by assuming that larger familiarity signals result from more vigorous input at the stage of familiarity encoding, and familiarity is reflected as repetition suppression (as we have observed). Under that assumption, the result will always be a representation in which familiarity and memorability intermingle in the population response vigor. As such, selective extraction of familiarity information will be required downstream to drive familiarity behavior without interference from memorability.

We observed this first type of intermingled representation of familiarity and memorability in ITC and we found evidence that the selective extraction of familiarity had been carried out in the MTL by examining HC. The most parsimonious explanation of these data is that familiarity encoding takes place within or before ITC, and the medial temporal lobe performs the selective extraction of familiarity, reflecting the results in HC. Because the model tested above is at the algorithmic level of analysis, it remains agnostic to the exact implementation of where these computations are implemented. Consequently, our main findings—that the transformation from familiarity happens via a two-stage process in which familiarity and memorability are initially intermingled and then familiarity is selectively extracted—holds whether the encoding initially takes place in ITC or elsewhere (it could in whole or in part be inherited or fed back from other structures), and it holds whether the extraction of familiarity is the result of a feed-forward transformation in MTL or happens in other ways, such as reciprocal computations between different parts of it.

### A simpler scheme cannot account for the transformation between ITC and HC

To recap, our results thus far suggest that a computational transformation occurs in the medial temporal lobe (MTL) between the ITC and HC representations. Linear decodability implies a weighted decoding scheme whereby some units are differentially weighted to create the downstream representation. Could this instead be achieved in an even simpler way, following on other ideas about hippocampal computation? Namely, visual representations in HC are more sparse than those in ITC^24,25^, a feature that could result from simple thresholding of incoming visual representations.

To address this possibility, we tested whether downstream decoding of ITC units with a threshold that chopped off the flanks of ITC tuning curves (thereby sparsifying their input) could account for the transformation observed in the neural data. For a mechanism like this to be plausible, familiarity information must be retained, but the modulation of firing by memorability should be reduced. In the linear decoding scheme presented in Fig. 3, this was the case: reading out ITC along a behaviorally aligned decoder axis did not cause catastrophic loss of familiarity information (Fig. S9).

To evaluate the plausibility of a simple “sparsification” mechanism, we began by fitting a model to each ITC unit to capture its visual tuning and familiarity sensitivity so we could simulate the effects of sparsifying ITC representations (Fig. 6a, see Methods). We confirmed that the model ITC population recapitulated salient aspects of our data, including the correlation between population response vigor and memorability (Fig. S15). Next, we applied the thresholding operation described in Fig. 6a, in which a percentage of the flank of each unit’s tuning function was set to a threshold, and explored over the full range of the tuning curve. The resulting model populations were analyzed for both their memorability correlations with firing vigor (Fig. 6b) and familiarity decoder performance (Fig. 6c). We found that matching the observed HC memorability correlation required thresholding a remarkable 93% of each unit’s ITC tuning function (Fig. 6b). The consequence of this amount of thresholding was the elimination of nearly all familiarity information (Fig. 6c). As such, simple thresholding (leading to sparsification) cannot account for the transformation from ITC to HC.

**Figure 6:**
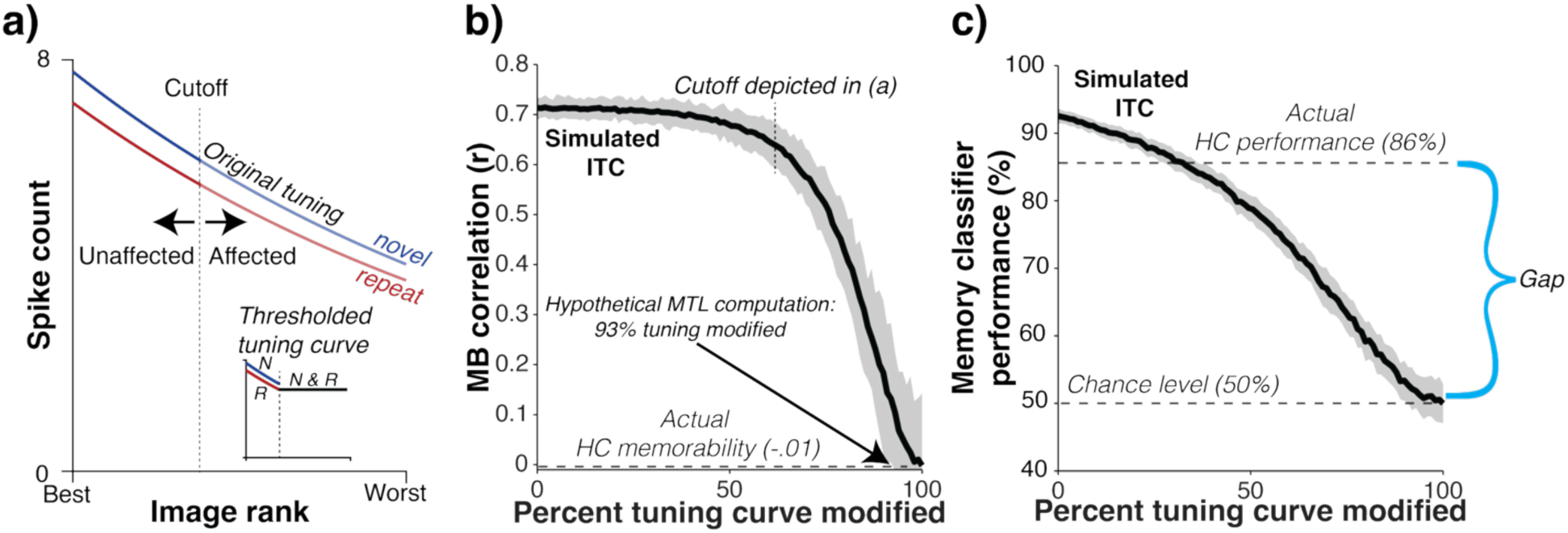
Thresholding ITC could not explain the transformation to HC. **a)** To evaluate the plausibility of the thresholding proposal, we performed a simulation based on tuning curves fit to each unit. Shown is the tuning for an example unit. The red and blue lines correspond to the firing of this unit to novel and repeated images, arranged from best (left, highest evoked firing) to worst (lowest evoked firing) images on the x-axis. The vertical dashed line denotes the threshold at which the tuning curve is modified, and the inset shows the tuning curve after the modification. To assess the impact of thresholding, the same threshold (e.g., 60% as shown in panel a) was applied to all ITC units, and the modified tuning curves were used to generate a simulated population; this was then repeated for a range of thresholds. **b)** The correlation between memorability and population vigor for the simulated populations, plotted as a function of the percent of the tuning curves that were modified on all units. The amount of the tuning curve modification can be thought of as the cutoff in Fig. 6a sweeping from right (0% modified) to left (100% modified). To match the observed memorability correlation in HC, at least 93% of the tuning curve would have to be thresholded away. **c)** Familiarity information remaining in each simulated population after thresholding, assessed by a weighted linear decoder. The threshold required to match memorability (highlighted by a cyan bracket) destroyed nearly all familiarity information, at odds with the reflection of robust familiarity information in the HC data (highlighted by a cyan bracket and the label “Gap”), implying that this proposal is implausible. Across the figure, the error shadows depict the 95% confidence interval. The same analysis windows were used as in Fig. 2-4.

## Discussion

Here, we provide evidence that a two-stage algorithm underlies the transformation from vision to familiarity, and we identify a previously undescribed medial temporal lobe computation that selectively extracts familiarity information from ITC. These results resolve the puzzle of how the brain differentiates between firing vigor modulations due to familiarity and memorability when determining image familiarity (Fig. 2) by demonstrating that a linear decoder can selectively extract familiarity information from ITC (Fig. 3). In addition to showing this could happen in principle, we find evidence that this extraction happens downstream of ITC with its outcome reflected in HC. There, memorability modulation is largely absent, but familiarity remains, and three specific predictions of this decoding scheme are confirmed (Fig. 4-5). Likewise, we’ve shown that a simpler scheme cannot account for the MTL transformation (Fig. 6), lending additional credence to our two-stage algorithmic description. That we observe differences in visual familiarity representations between ITC and HC is notable in light of recent work in monkeys showing that ITC and HC representations largely align in a familiarity task in which they are expected to be different^16^ (namely, familiarity for visually-similar images, which has been tied to the hippocampus via fMRI, modeling, and natural lesion studies^26–30^).

These results lead to a simple, two stage algorithmic account of how seeing an image is transformed into a memory that it has been seen. The process begins with the retinal input, where firing vigor is not modulated by either memorability or familiarity. In Stage 1, which occurs across the ventral visual pathway, more memorable images come to produce more vigorous responses (Fig. 7, middle). This, in turn, causes more repetition suppression. Under the assumption that repetition suppression is a proportional reduction in the stimulus-evoked response, the more vigorous responses associated with more memorable images lead to larger reductions (e.g., 10% of 10 leads to a reduction of 1; 10% of 50 leads to a reduction of 5, Fig. S3). However, this leaves an ITC representation in which familiarity and memorability intermingle. We found that this representation can be linearly decoded to extract familiarity information despite memorability modulations potentially interfering with readout of RS (Fig. 3). In Stage 2, familiarity is selectively extracted (cyan arrow in Fig. 7, middle) while leaving behind memorability such that a largely isolated familiarity representation is reflected in HC (Fig. 7, right). This is consistent with the predictions of the linear readout of ITC (Fig. 4-5). We infer that this representational transformation took place between the two brain areas, in the medial temporal lobe.

**Figure 7:**
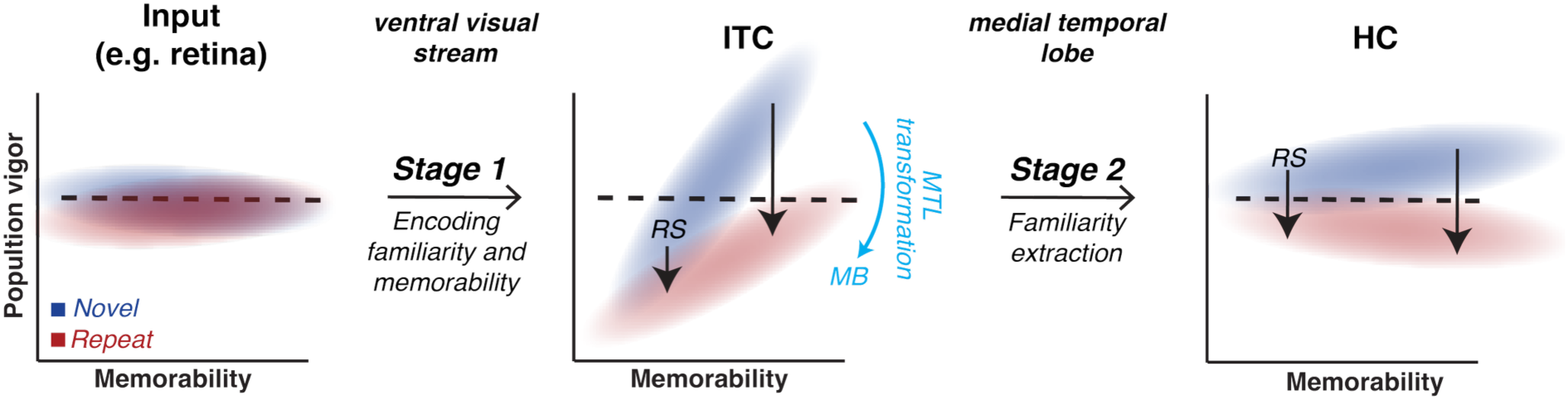
Schematic of the two-stage algorithmic transformation creating the ITC and HC representations. In each of the plots, the blue cloud represents a cartoonized distribution of firing vigor to novel images and the red cloud to repeated images. Left: Early in the visual system, such as in the retina, memorability and familiarity are not reflected in firing vigor. In Stage 1 of the algorithmic transformation, the ventral visual pathway (e.g. retina, LGN, V1, V2, V4, and IT) creates an ITC representation (middle), where both memorability (MB) and repetition suppression (RS, black arrows) influence the firing vigor. Namely, more memorable images elicit stronger responses, resulting in greater repetition suppression. However, this results in an ITC representation in which both memorability and familiarity intermingle. In Stage 2 of the algorithmic transformation, familiarity is selectively extracted, amounting to a rotation of the distributions shown in ITC (cyan arrow) to create the HC representation. In HC, familiarity is reflected largely independent of memorability.

Although this account relies on a comparison between ITC and HC representations, the implementation of Stage 2 may occur in one or more MTL structures (such as perirhinal cortex), may arise gradually across the MTL, or singularly within HC, and any one or all of those brain areas may ultimately contribute to memorability behavior. That is, the model proposed here holds explanatory power at the level of algorithm. Under the assumption that larger familiarity signals result from more vigorous input at the stage of familiarity encoding, coupled with the assumption that familiarity is reflected as repetition suppression (both reflected in ITC data), the result will always be a representation in which familiarity and memorability intermingle in population response vigor. As such, selective extraction of familiarity information will be required downstream to drive familiarity behavior without interference from memorability. Our model describes how this transformation happens algorithmically.

This account suggests the HC familiarity information is inherited from earlier areas. While future causal manipulations would be required to definitively map these circuits, the data and literature support this account. First, the familiarity signals observed in HC mirror the visually specific format and repetition suppression present in the ventral visual stream, aligning with the predictions of the behaviorally aligned readout of ITC (Figs 4-5). Second, evidence from lesion studies in humans^31^ and animals^32^ shows that damage to HC leaves overall familiarity performance intact in scenarios like the one we test here, suggesting that HC is not the sole locus or origin of familiarity storage. These factors present issues for any alternative class of models in which the neural signals that drive familiarity originate in HC (not ITC).

Given the finding that memorability behavioral patterns are linearly decodable from ITC, why might the brain bother to remove variation caused by visual features (like memorability) downstream? Why not just retain them in a linearly decodable format? Though impossible to know, we can speculate. Whereas “pattern-of-spikes” coding schemes (where some neurons fire more than others to convey information) are associated with variables such as the identities of the objects in the world, “magnitude” coding schemes (where variables are reflected in the vigor of a population or subpopulation) have been associated with the degree to which that information should be prioritized^33^. Once familiarity information is encoded, eliminating memorability modulation could free up magnitude coding bandwidth for other priority-related variables, such as those associated with reward, attention, and prediction. Future studies manipulating multiple priority codes would be required to fully assess this possibility.

The hypothesis that the MTL carries out the transformation described between ITC and HC predicts that damage to the MTL, where we propose that the familiarity extraction computation happens, should lead to altered patterns of memorability in patients with mild cognitive impairment (MCI, which is associated with MTL atrophy^34^) compared to healthy controls. Relevant to that prediction, one study^35^ investigated memorability in individuals with MCI and found that memorability scores were less predictive of behavioral performance in patients with MCI than for healthy controls, though there was still some overlap in images that both healthy controls and MCI patients found memorable. However, it’s unclear whether this resulted from the specific memorability changes our results predict or from overall reduced memory performance, leading to lower inter-subject consistency in these patients; thus, further investigation is warranted. The model we proposed above makes the testable prediction that if the MTL is responsible for extracting familiarity from the visual cortex, reduced MTL function should systematically reverse memorability’s effect on behavior (like Fig. 2b) in patients with MTL damage. If true, testing for this could be a non-invasive screening method for MTL damage or degradation.

Future work will be required to determine how the MTL might learn to decode ITC in this way. The memorability decoder used in Fig. 3 and 5 was trained with knowledge of the memorability of each image, in question is how the brain might have access to that variable or a proxy for it. The content of images can be predictive of memorability^36^, creating a defined statistical landscape^37^ that the brain could, in principle, learn for this purpose. Underscoring this point, neural networks trained to predict memorability are able to do so for unseen images^5,38^, a tool utilized in this study to generate memorability scores for tens of thousands of images, but also a demonstration that memorability has a learnable statistical structure, and the brain could learn these statistics as well. Likewise, for the decoding scheme described in Fig. 3 to work, there must be meaningful heterogeneity in the sensitivities of individual ITC units to familiarity and memorability (or their representations would completely overlap and the angle difference between their decoding axes would be zero). That the decoding scheme implemented in Fig. 3 finds these axes are separated in ITC and faithfully maps responses to behavior reveals that this heterogeneity exists in ITC and suggests that the MTL could learn these differences to selectively extract familiarity information.

The structure of the behavioral task used here means that these results specifically pertain to how the brain supports familiarity for images seen exactly once before. Therefore, these results are not directly comparable to those derived from paradigms involving extensive experience following hundreds or thousands of image encounters over the course of days to weeks (such as refs^39–42^). For instance, while we observe that single-exposure familiarity is consistently reflected as repetition suppression in both ITC and HC, studies utilizing hundreds of exposures to highly familiarized objects have linked ‘old’ judgments to repetition enhancement (rather than suppression) in other medial temporal lobe structures such as the perirhinal cortex^40^. These contrasting signatures may reflect distinct neural mechanisms required for single-exposure familiarity vs highly trained familiarization.

While the simulations shown in Fig. 6 show that a thresholded readout (something that could generate the sparsity supporting decorrelation and pattern separation in HC) cannot by itself lead to attenuated memorability representations, our results may relate to other proposed schemes for HC. For example, hippocampal place cell-like representations can arise via an autoencoder-based scheme designed to compress memories^43^, and the architecture of hippocampus has been linked to learning statistical regularities in the environment^44^. The attenuation of memorability observed in HC relative to ITC could conceivably be linked to these or other computations and should serve as a testable constraint on future computational models of visual familiarity processing by the medial temporal lobe.

Understanding how the healthy brain transforms experiences into memories is foundational for understanding memory-related disorders. By focusing on the nature of the neural code that the brain uses to drive familiarity reports, we uncover a previously undescribed computation that can constrain future models of medial temporal lobe function and may help make sense of the behavioral patterns of individuals with medial temporal lobe damage. In addition, we present a more complete understanding of how seeing an image is transformed into familiarity that can be tested and refined in the broader pursuit of understanding how memory and its underlying signals come to be, both in health and disease.

## Methods

All data were collected from two adult rhesus macaque monkeys (*Macaca mulatta*). Monkey 1 was male and monkey 2 was female. All procedures were performed in accordance with the guidelines of the University of Pennsylvania Institutional Animal Care and Use Committee.

### Anatomy and recording

The activity of neurons in ITC and HC were recorded via a single recording chamber (Crist Instruments, Hagerstown, MD) in each monkey. In monkey 1, we recorded from left ITC and HC. In monkey 2, we recorded from right ITC and HC. Chamber placement was guided based on anatomical magnetic resonance imaging. ITC and HC were recorded in separate sessions. Neural signals were recorded with 24-channel U-probes (Plexon Inc., Dallas, TX) with recording sites spaced at 100 μm intervals. Wideband signals were amplified and digitized at 30 kHz using a Grapevine Data Acquisition System (Ripple, Inc., Salt Lake City, UT). Spikes were hand-sorted offline (Plexon Offline Sorter).

The region of ITC recorded was located on the ventral surface of temporal lobe in anterior ITC where repetition suppression in response to familiarity has been previously observed^11^; this region is similar to that recorded in ref^15^.

M1’s ITC population was recorded from four locations ranging from 17-19mm lateral to the sagittal plane defined by the midline (approximately 1-3mm medial to the anterior middle temporal sulcus (amts)) and 14-15mm anterior to the coronal plane defined by the ear canals.

M2’s ITC population was recorded from four locations ranging from 17-18mm lateral to the sagittal plane defined by the midline (approximately 0-1mm medial to the AMTS) and 13-14mm anterior to the coronal plane defined by the ear canals.

The region of HC recorded only included areas in the dorsal half of HC (this targeting was guided by ref^22^). This means that the sample predominantly includes hippocampal subfields CA3, DG and CA4 (approximately in that order of relative prevalence) but because of the organization of the primate hippocampus and the inherent ∼+/− 1 mm level of precision in our recording preparation, we do not attempt to distinguish between these subfields. However, because subfield CA1 is located ventrally in primate hippocampus and we targeted our recordings to the dorsal half of HC, we are reasonably confident it is not represented in these populations.

M1’s HC population was recorded from seven locations ranging from 13-15mm lateral to the sagittal plane defined by the midline and 14-16mm from the coronal plane defined by the ear canals.

M2’s HC population was recorded from 6 locations ranging 10mm-12mm lateral to the sagittal plane defined by the midline and 11-13mm anterior to the coronal plane defined by the ear canals.

We did not observe any obvious functional organization within the regions recorded in ITC or HC.

### Inclusion Criteria

A multi-channel recording session was included in the analysis if (1) there were no technical problems with the recording hardware, (2) it fell within the anatomical boundaries described above, (3) the monkey completed at least 200 novel-repeat image-pairs during the session, (4) the session was stable, defined as the overall baseline firing rates across all channels did not change by more than 2-fold across the session, and there was no significant difference between baseline firing rates preceding novel and repeated images (t-test, p<.01, in the 500ms prior to image presentation) and (5) the session was composed of over 50% visually responsive units (in ITC) or contained at least a single visually responsive unit (HC), determined by an inspection of the raster plots. We chose this relatively strict criterion for ITC to ensure high-quality data based on our previous experience recording this highly visual brain area (based on extensive previous experience, a lack of visually responsive units in a nominally ITC session suggested the probe was actually located in white matter dorsal to ITC), and a much looser criterion for HC because we did not wish to make any *a priori* assumptions about its visual organization. Approximate familiarity information was matched between populations by recording HC data in monkey 2 until FLD familiarity decoder performance in the 200-500 window matched that of the ITC population, and simulating this procedure by truncating a handful of ITC sessions in monkey 1 (criterion 6). Within a session, units were included in the population if they evoked a response in the post-stimulus window [50-350ms] versus the pre-stimulus [-300,0] window (two-sided t-test, p<0.10) across all trials in the session. In monkey 1, this filter passed 96% and 52% of units for ITC and HC respectively, and for monkey 2 this filter passed 93% and 57% of units for ITC and HC respectively.

Monkey 1’s ITC population was collected over the course of approximately five weeks. We collected 20 sessions and 13 were removed (1, 1, 1, 6, and 4 from criteria 1, 3, 4, 5 and 6 respectively).

Monkey 1’s HC population was collected intermittently over the course of approximately seven months. We collected 36 sessions and 20 were removed (3, 8, 3, and 6 from criteria 1, 3, 4 and 5 respectively).

Monkey 2’s ITC population was collected intermittently over the course of approximately four months. We collected 9 sessions and 1 was removed (from criteria 1).

Monkey 2’s HC population was collected over the course of approximately two months. We collected 30 sessions and 9 were removed (2, 3, 2, 1, and 1 from criteria 1, 2, 3, 4, and 5 respectively).

### Behavioral task

A custom MWorks script (The MWorks Project, Cambridge, MA) was used to display images on an LCD monitor and collect behavioral responses. The experiment was performed in a darkened testing chamber with the monkeys’ heads fixed while their gaze was tracked with an Eyelink 1000 (SR Research, Ottawa, Canada). Trials were initiated by the monkey fixating on a red square fixation point (0.35 degrees) in the center of a gray screen within an invisible square window (4 degrees monkey 1, 3.5 degrees monkey 2), followed by a 500ms delay before a 4-degree image appeared at the center of the screen. The monkeys had to maintain fixation on the fixation point for 500ms, after which the image disappeared from the screen and the fixation point changed to a green go cue, and two square white 0.5 degree response targets appeared 8 degrees above and below the stimulus. The monkey indicated a response by making a saccade to one of the two targets. If the monkey was correct, they received a liquid reward. For Monkey 1, the target associated with “novel” images was the top target; this was reversed for Monkey 2. If the monkey broke fixation, the image immediately disappeared, and a new trial was offered after a short delay. In cases where the stimulus was novel but the monkey broke fixation (after an image had appeared on the screen), the subsequent presentation of that image was still rewarded as a “repeat,” although incomplete image pairs were excluded from further analysis.

A custom MATLAB (The MathWorks, Inc., Natick, MA) script was used to pseudo-randomly generate a sequence of images, where each image appeared exactly twice (the same algorithm as ref^15^). The script aimed to match a target distribution of n-back trials. To fill the sequence, some occasional off-target n-backs were used but were not included in analysis. Due to behavioral differences between the two monkeys, the exact n-back distributions varied between Monkey 1 and Monkey 2.

Monkey 1 was presented with n-back trials of 1, 2, 4, 8, 32, 48, 64, and 192. The distribution was uniform for n-backs between 2 and 64, while 1 and 192 were half as frequent (e.g., in a sequence, there could be 15 1-back trials, 30 2-back trials, 30 4-back trials, and so on).

Monkey 2’s sequence included n-backs of 1, 2, 4, 8, 16, 32, 48, and 64, with a uniform distribution between 2 and 48 and half-frequencies for 1 and 64. N-backs differed between monkeys in order to approximately match performance between animals.

The images used were drawn from the same set used in Jaegle et al. (2019)^8^, and images used during training a monkey were never reused during recording.

### Pseudopopulation

Because the responses of individual ITC and HC units are often tuned for specific visual features, estimating the overall population response requires many hundreds of units. Therefore, we concatenated units across sessions into a pseudopopulation and aligned images of similar memorability into “pseudoimages”. In the pseudopopulation, a response to a given pseudoimage pair always contained the novel/repeated pairing of the same real images, and pairs were aligned so that a pseudoimage always included a similar n-back separation and images with similar memorability scores. We also used these alignments to pool the data between monkeys. Because sessions differed in numbers of completed novel/repeat image-pairs, the final size of the pseudopopulation was limited by the session with the fewest number of completed pairs. To align sessions, in the longer sessions, images were ranked and uniformly subsampled by memorability to obtain a similar distribution of memorability across all sessions, as in ref^8^ before aligning and concatenating.

### Vigor decoder (Fig. 3 and 5)

This decoder was cross-validated and took the form of a linear discriminator in which the class of a population response vector to a pseudoimage was determined by the sign of:

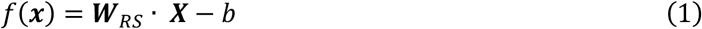

Where ***X*** is the mean-centered matrix of spike counts to each unit and pseudoimage, and a value of f(x) > 0 classifies an image as novel, and f(x) < 0 as repeated. **W** evenly and positively weights each unit so that:

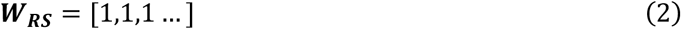

And:

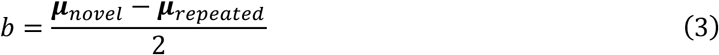

### FLD decoder (Fig. S5 and S11)

For the FLD decoder, we used the same implementation as refs^15,17^. The decoder was cross-validated with 1000 splits, and in each split 70% of the data were used for training and 30% for testing. The decoder took the form of a linear discriminator in which the class of a population response vector to a pseudoimage was determined by the sign of:

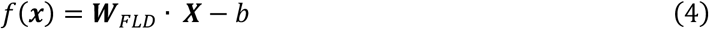

Where **X** is the population response vector, ***W****_FLD_* is an n-dimensional weights vector in the n-dimensional ITC or HC neural space (n being the number of units), and b is a decision boundary given by:

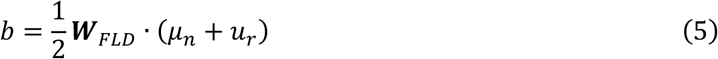

Where *μ_n_* and *u_r_* are the means to the two classes (novel and repeated) and the weights are calculated by.

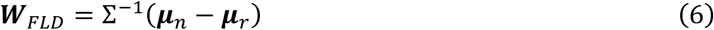

Where Σ^-1^ is the inverse of the covariance matrix averaged across both conditions. Because the dimensionality of the neural data is very high relative to our sample size, we cannot make good estimates of the off-diagonal terms in the covariance matrix so they are set to zero. This results in the decoder effectively weighting each unit by its d’.

### Ranked FLD (Fig. S11)

The same implementation of the FLD decoder described above was used. Weights for all units were calculated within each cross-validation iteration, then units were ranked by those weights, subsets of units were removed, and the decision boundary was re-calculated before testing the decoder. Because we were interested in comparing ITC and HC, and as documented in Fig. S5, memorability confuses decoders of this type, for Fig. S11, the pseudopopulation alignment by memorability was broken by randomly shuffling the images that compose pseudoimages. This means that every pseudoimage was composed of random images, ameliorating the counterintuitive influence of memorability and allowing comparison of total familiarity information between brain regions.

### Memorability and behaviorally aligned decoders (Figs. 3 and 5)

To train and test a decoder that might account for the transformation between ITC and HC, we used similar methods to ref^17^. The analysis was cross-validated with 1000 splits, and in each split 60% of the data was used for training the MB decoder, 20% to find the location of the behaviorally aligned axis and 20% held out to test behavior at the behaviorally aligned axis.

The vigor decoder uniformly weighted each unit equally and positively, as described above in equations 1-3.

The MB decoder was similar to the vigor decoder, but the weights were defined by:

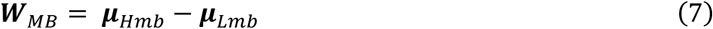

Where ***μ****_Hmb_* is the vector of unit’s mean response to images in the top 50% of memorability scores and ***μ****_Lmb_* is the mean responses to images in the bottom 50% of memorability scores.

The decoders rotated within the plane took the form of:

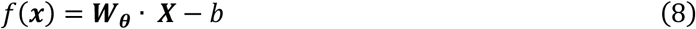

Where **X** is the mean-centered matrix of spike counts to each unit and pseudoimage projected onto the MB/RS plane and where W for an angle *θ* on the MB/RS plane is defined by:

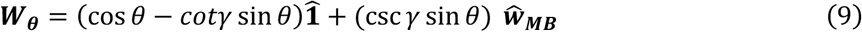

Where **1̂** is the unit vector representing the RS axis, and ***ŵ_MB_*** is the unit vector representing the MB axis and *γ* is the angle between the MB and RS axes. And

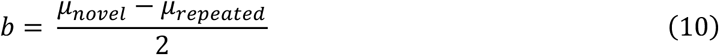

### Behavioral predictions based on decoders (Figs 2b, 3b, 5b and S5)

Any comparison between population decoded predictions and actual behavior requires implicitly or explicitly specifying the number of neurons included in the decoding analysis. Following on ref^17^, we confirmed that performance using all recorded units in our dataset fell below saturation, and then simulated increases in population size by fitting a single rescaling parameter by solving for a single multiplicative scaling factor that minimizes the squared error between the neural predictions and behavior, and then multiplied the predictions by this rescaling factor. This approach is designed to match overall performance while respecting the relative relationship between points. In cross-validated analyses, rescaling took place once after averaging across all runs. Any rescaled predictions over 100% were clamped to 100.

### Projections onto plane defined by 1̂ and *ŵ_MB_* and error ellipses (Figs. 3a and 5a)

The data used to visualize the contour ellipses were the fully held-out test set, split into four divisions by memorability quartile boundaries. The ellipses were computed by projecting the data onto the non-orthogonal **1̂** and ***ŵ_MB_*** axes. This coordinate system was then rotated by 45 deg to visualize the RS decoder at this angle. For each condition, the covariance matrix of transformed data was computed, and eigenvectors of this matrix provided the major and minor axes of the associated ellipse. The ellipses themselves represent one standard deviation contours of the 2-dimensional histograms projected onto this plane. The data plotted in Fig. 3a and 5a represent the average across all cross-validation iterations.

### Prediction quality and behaviorally-aligned axis (Figs. 3 and 5)

To determine the quality of the prediction between actual behavior and that predicted by decoded performance, we divided the testing set for each round of cross-validation into four memorability bins with the quartile boundaries defined by the quartiles of the overall memorability distribution (these were also used to plot the contours, see above). They were further divided by novel and repeated images, and novel and repeated prediction qualities were calculated separately. These sets were used to create decoded predictions of behavior. To quantify prediction quality (PQ), we fit a linear regression to both the behavioral and neural predicted data and compared the slopes of the regressions such that:

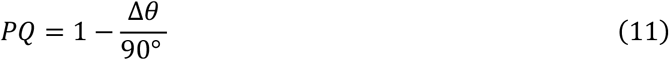

Where *θ* is the angle between the behavioral and neural-predicted slopes with a possible range of 0 to 90 degrees. This metric yields a value of 1 when the slopes are perfectly aligned and 0 when they are perpendicular. The final PQ used to determine ‘behaviorally aligned’ decoder axis is the mean of this result for novel and repeated PQ.

### Behavioral predictions on the behaviorally aligned axis (Fig. 3b and 5b)

Within each cross-validation split, a behaviorally aligned axis was identified. To make behavioral predictions, in each run, the fully held out data (not used to find the MB decoder or the best-PQ axis) was projected onto that axis, yielding a PQ and a behavioral prediction. The final plotted behavior is the average across cross-validation runs.

### Estimates of error in decoder results

Consistent with other cross-validation procedures^45^, the standard error was computed as the standard deviation of the mean across 1000 cross-validation iterations.

### Bootstrap analysis for difference in PQ (Fig. 3 and S6-S7)

To test whether prediction quality (PQ) significantly differed between the BA and VG axes, we used a bootstrap analysis, resampling across neural units. For each bootstrap sample, units were sampled with replacement from the population and the full cross-validation procedure was repeated on this resampled population. This produced bootstrap distributions of test-PQ for the BA and VG axes. Statistical significance was assessed from these distributions as the number of bootstrap differences (out of 1000) that differed from the null hypothesis (of no difference between BA and VG axes); if none differed, the p-value was reported as <1e-4. The 95% confidence intervals were computed from the 2.5^th^ and 97.5^th^ percentiles in the bootstrap distribution.

### Permutation test for difference in slopes (Figs. 4 and S13)

To perform this test, a null hypothesis distribution was generated by randomly assigning an “ITC” or “HC” label to each data point, without replacement, then fitting lines to these shuffled “populations” and calculating the difference in slopes between the two “brain areas”. This procedure was repeated 10,000 times to generate a distribution of the null hypothesis that there was no difference between the brain areas. P-values were calculated by computing the proportion of null-distribution differences in slopes that differed from the actual slope.

### Repetition effect (d’ for familiarity, Fig. 4)

d’ for a unit was calculated by:

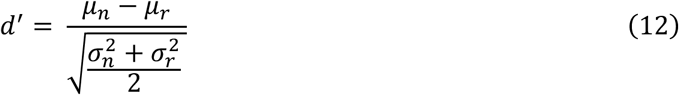

Where *μ_n_* and *μ_r_* are the mean to novel and repeated images, respectively, for that unit, and 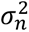 and 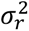 are the variance within novel and repeated images, respectively, for that unit.

### Population fits/synthetic population generation and modification (Fig. 6)

We fit units following an adaptation of the method described by ref^17^. In brief, this involved fitting a three-parameter exponential model for each unit that maximized the likelihood of observing the spike count data from 100-500ms:

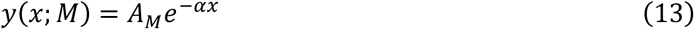

Where *x* is the stimulus rank along the tuning curve, M is the familiarity condition (novel or repeated), A is amplitude (*A_N_* and *A_R_*, for novel and repeated images, respectively, fit separately), and *α* controls the sharpness of the curve (to capture image selectivity).

To generate synthetic populations based on our observed data, we ranked the observed spike counts for each image seen (averaging across novel and repeated presentations) to get an *x* value (image rank) for each image for each unit. These values were then used to solve for a predicted spike count for a given image and unit using the fit constants for *A_N_*, *A_R_* and *α*. We then used a Poisson process to generate a synthetic spike count. Memorability in these simulations derives from the memorability scores of the originally shown images

To generate the ‘modified populations’ in Fig. 6, the predicted firing rate for both novel and repeated was changed to that of the repeated firing for any image ranked lower on the tuning curve than the ‘cutoff point’ rank.

## Acknowledgements

This work was supported by the National Eye Institute of the NIH (award R01EY020851 to N.C.R.), the Simons Foundation (Simons Collaboration on the Global Brain award 543033 to N.C.R.), and the National Science Foundation (award 2043255 to N.C.R.).

## Supplementary Figures

**Figure S1:**
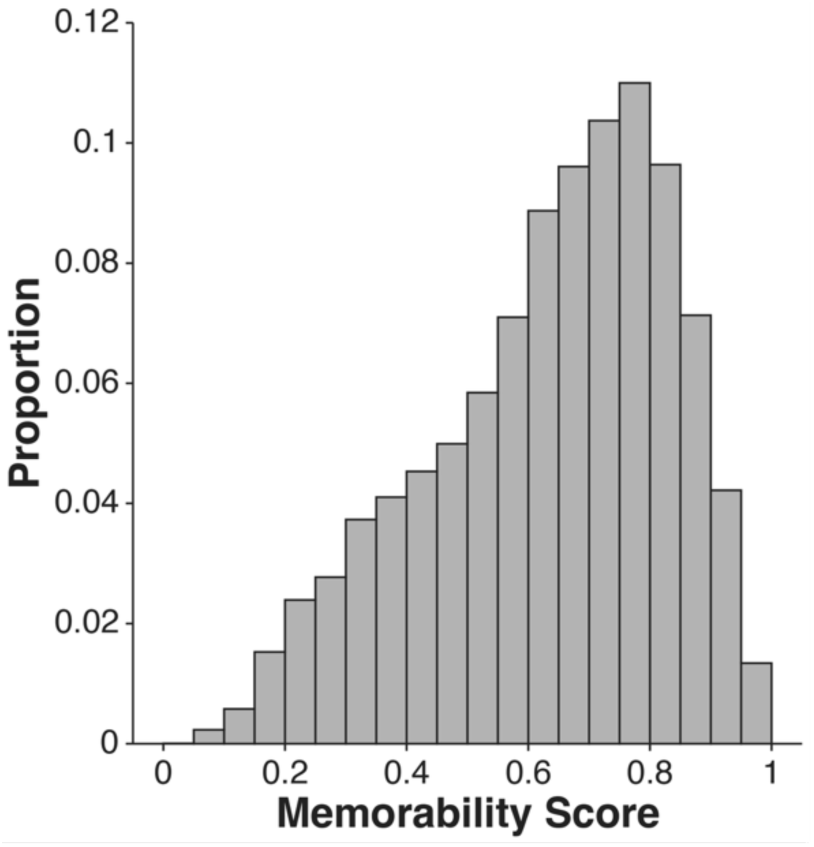
Distribution of memorability scores across all images. Images (n=10,816 images where both novel and repeated trial were completed) were scraped from the internet and covered a wide variety of objects and scenes. Memorability scores were acquired using the convolutional network MemNet^5^, which is trained to predict human visual recognition memory on a similar familiarity task.

**Figure S2:**
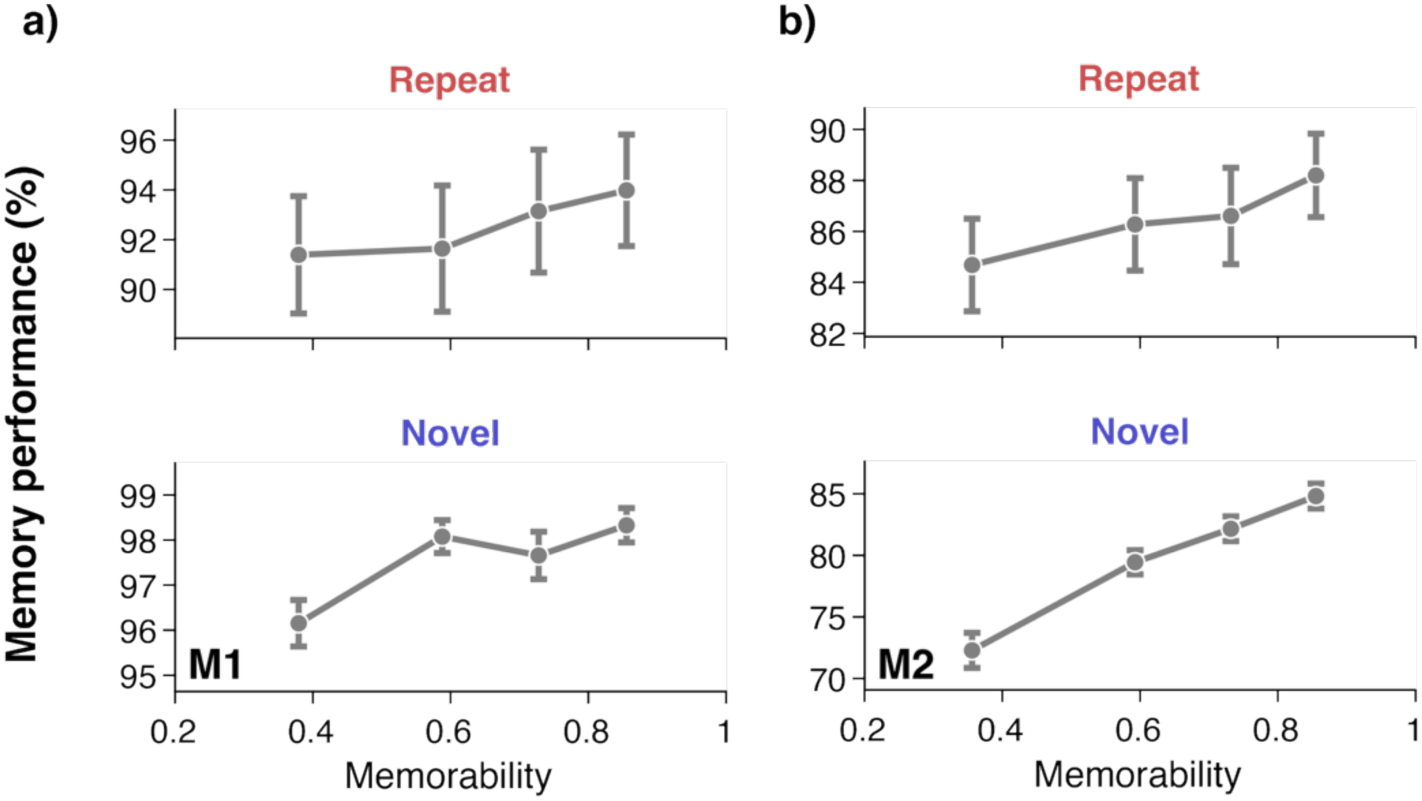
Individual monkey behavior. Gray dots show means of four memorability bins of equal size, error bars are the standard error of the mean and are larger for repeated than novel because repeat encompasses a range of n-backs but all novel trials are the same. **a)** In M1, there were 4,784 trials for each condition (novel and repeat). There was a significant effect of memorability on novel and repeated behavior (logistic regression on repeated trials *β* = 1.19, odds ratio = 3.27, p = 9.19e-6. For novel trials *β* = 1.49, odds ratio = 4.44, p = 8.7e-4). **b)** For M2, there were 6,032 trials for each condition and memorability also significantly affected behavior (*β* = 0.62, odds ratio = 1.86, p= 4.9e-4. For novel trials, *β* = 1.58, odds ratio = 4.87, p = 9.80e-4).

**Figure S3:**
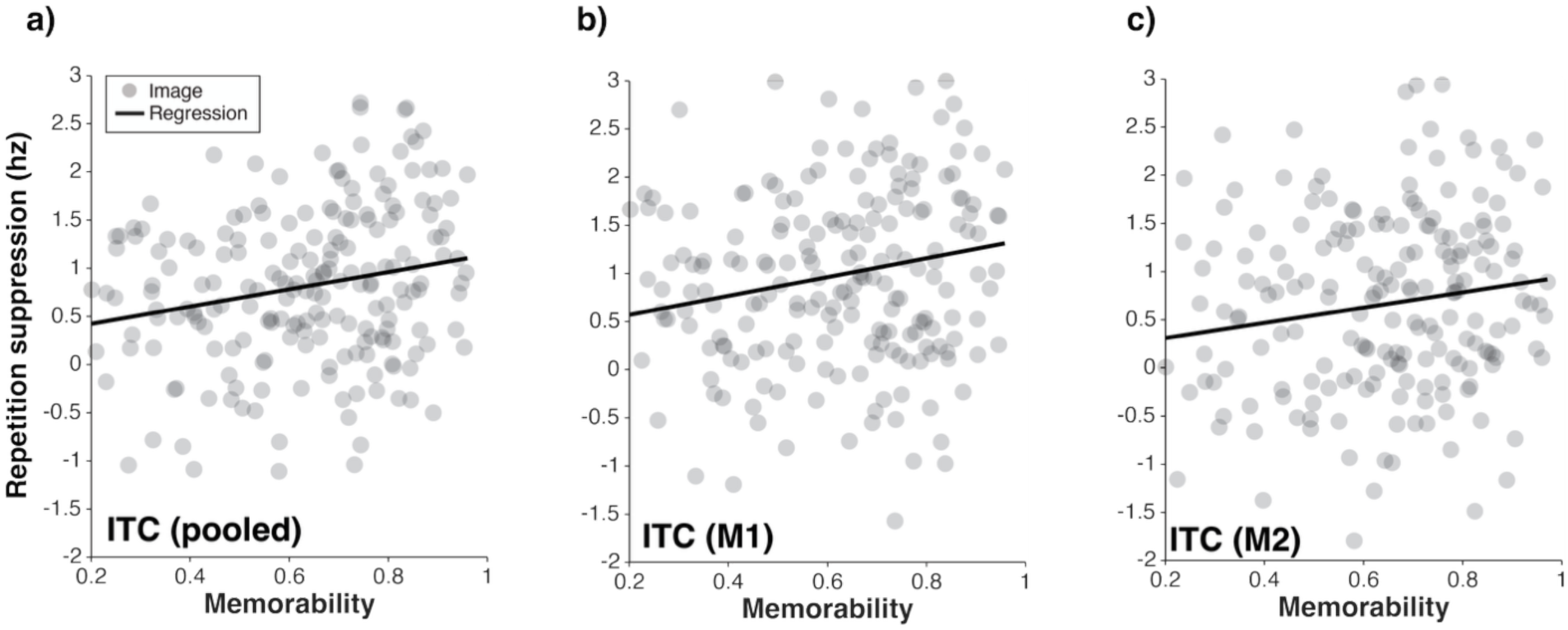
Repetition suppression (RS) as a function of memorability in ITC. Each dot represents RS for a given image, computed as the difference in firing rate across all units to the novel versus repeated presentation of that image with a given memorability; lines depict linear regressions. **a)** There was a significant correlation between memorability and RS in the pooled population (r(206) = 0.23, p = 0.001). **b)** In M1, r(206) = 0.21, p = 0.003, **c)** in M2, r(206) = .15, p = 0.028. Data were analyzed in the 100-500ms window following stimulus presentation.

**Figure S4:**
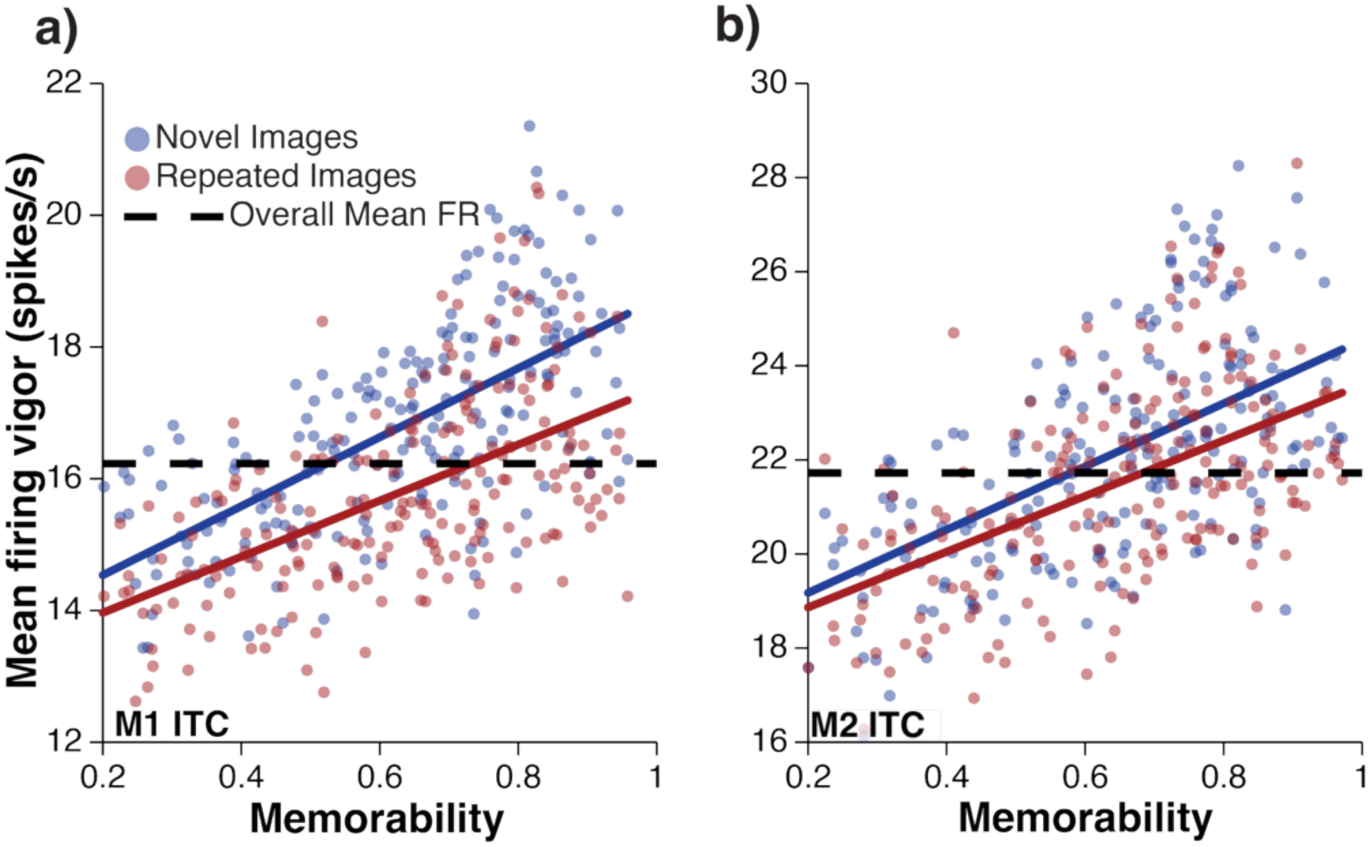
Relationship between memorability and firing rate in each monkey analyzed individually. Same convention and analysis windows as Fig. 2a **a)** In M1, there was a significant correlation between memorability and firing vigor for repeated images (r(206) = .58, p = 1.8e-20) and novel images (r(206) = 0.67, p = 2.1e-28) **b)** In M2’s ITC, there was a significant correlation between memorability and vigor for repeated images (r(206) = 0.56, p = 1.0e-18) and for novel images (r(206) = 0.60, p = 2.1e-21).

**Figure S5:**
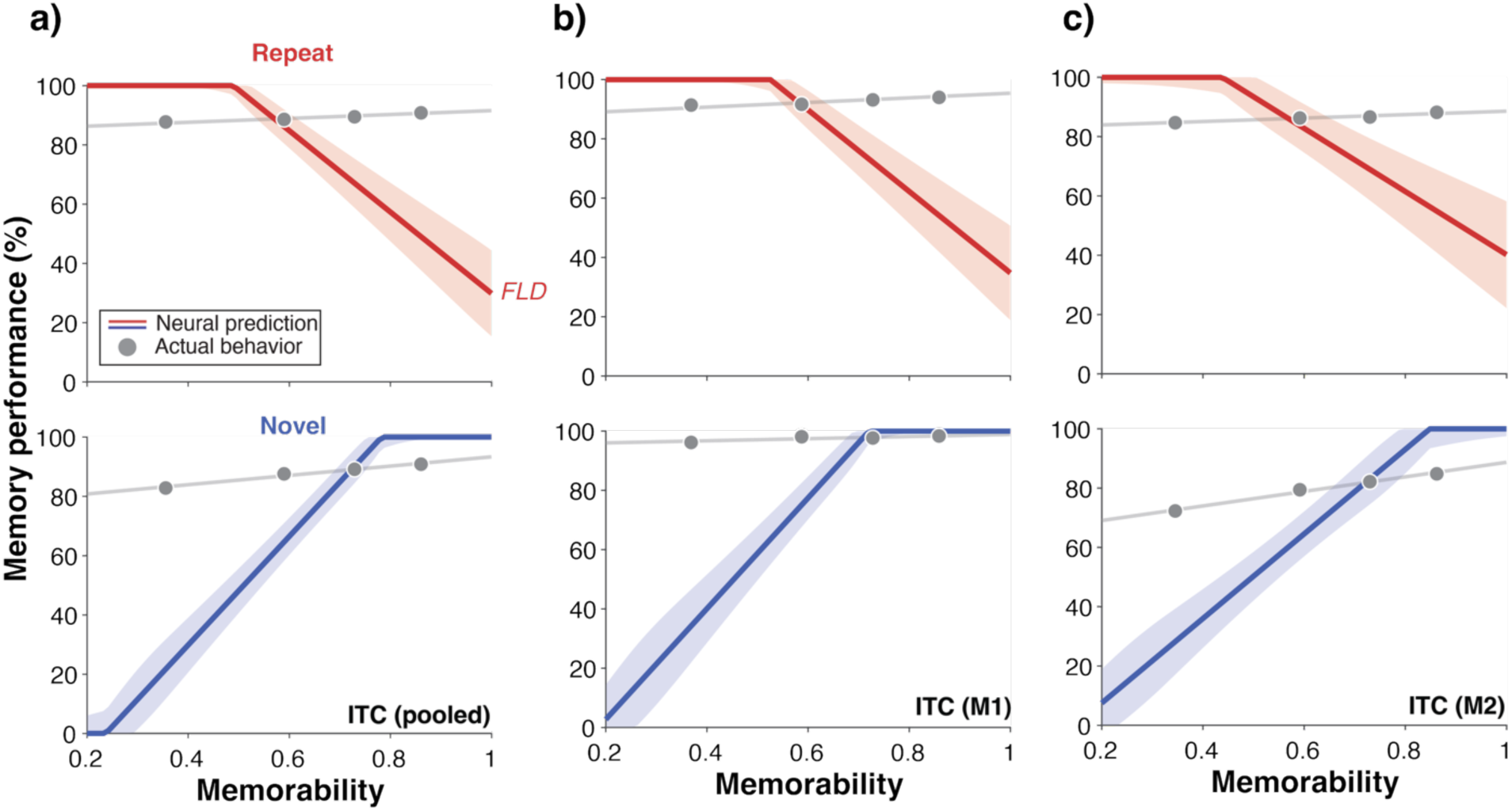
Similar to decoding population vigor alone, an optimized weighted linear decoder also produced similarly misaligned predictions of memorability behavior. Plotted here are behavioral predictions resulting from training a Fisher’s linear discriminant decoder (FLD, see Methods) on all neural spiking to images of all memorability scores and then testing it on binned memorability subsets, as in Fig. 2b. This decoder differed from the vigor (VG) decoder tested there by being able to weight each unit by its informativeness and incorporate any units that might be enhanced by repetition with a negative weight. The results were predictions that were misaligned in the same way as the VG decoder. **a)** In the pooled data, the mean +/− se prediction quality (PQ, a measure from 0 to 1 quantifying alignment between the predicted and actual behavior, see Methods) was 0.42 +/− 0.07 for repeated images and 0.46 +/− 0.04 for novel images. **b)** In M1, the PQ for repeated images was 0.43 +/− 0.07 and 0.42 +/− 0.06 for novel images. **c)** In M2, the PQ for repeated images was 0.55 +/− 0.11 and 0.62 +/− 0.08 for novel images. Data were analyzed in the 100-500ms window following stimulus presentation. Raw decoder outputs were scaled using a single multiplicative scaling factor following the same procedure as Figs 2, 3, and 5 (see Methods).

**Figure S6:**
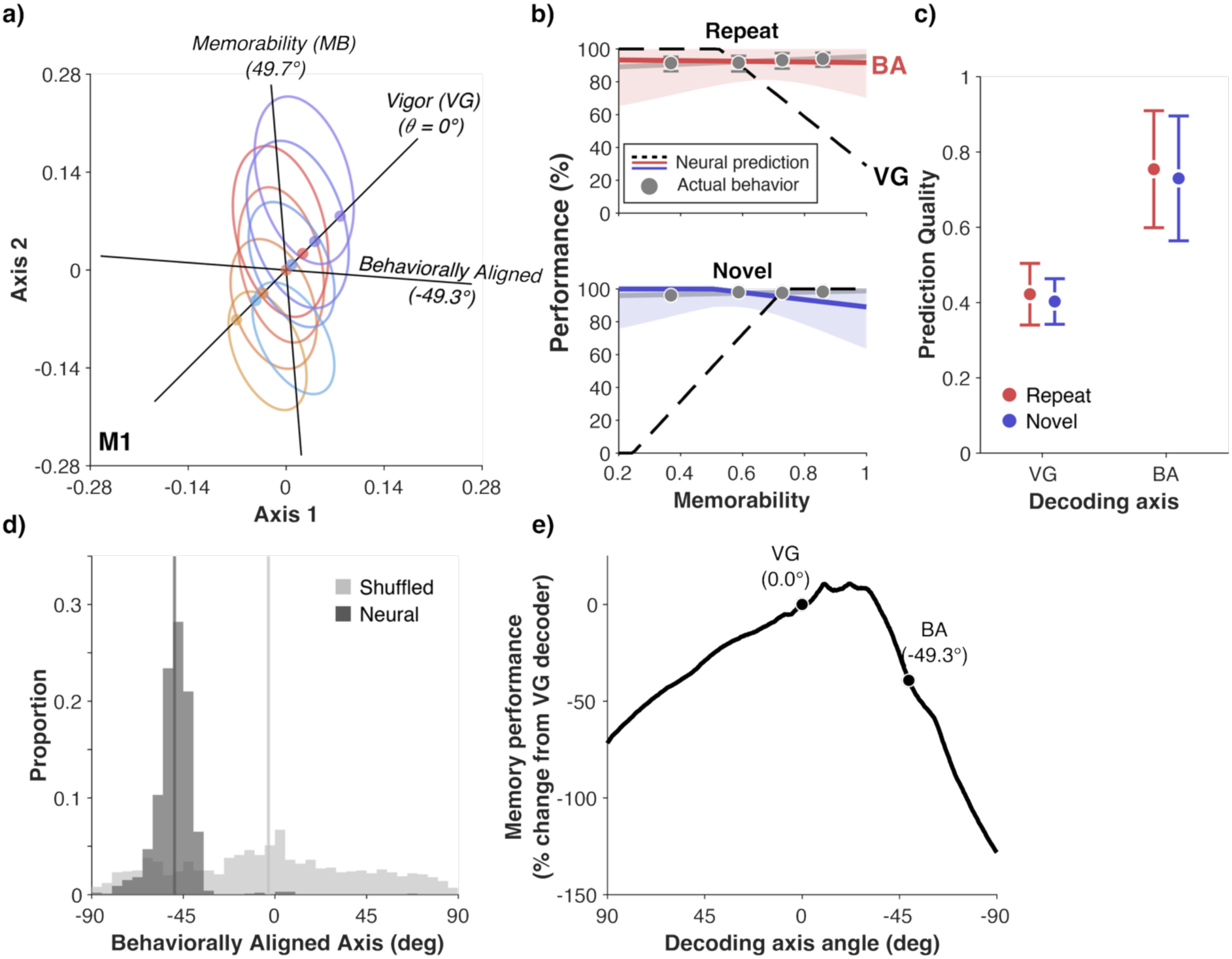
Memorability behavior is linearly decodable from ITC (M1). Same conventions as Figs 3 (top row) and S9 (bottom row). **a)** Projections of ITC neural data onto a plane defined by an axis corresponding to vigor decoder (where every unit is weighted equally, ‘Vigor (VG)’) and a linear memorability decoding axis optimized to decode high versus low memorability images, ‘Memorability (MB)’. **b)** Comparison between observed behavior (gray) and neural predictions for the vigor decoder axis (dashed line) and the behaviorally aligned decoding axis (BA, red and blue lines correspond to repeated and novel behavior, respectively). **c)** Comparison between prediction quality when decoding behavior using the vigor (VG) axis versus the behaviorally aligned (BA) axis using the same fully held-out data as described above. Decoding along the VG axis produced poor predictions of memorability behavior (repeated PQ mean +/− se: 0.42 +/− 0.08, novel PQ: 0.40 +/− 0.06). Decoding along the BA axis produced good predictions of both repeated and novel behavior when tested on fully held-out data that was not used to train either the memorability decoder or to find the BA axis (repeated-behavior PQ mean +/− se: 0.75 +/− 0.16, novel: 0.73 +/− 0.17). A bootstrap analysis (see Methods) revealed that the BA decoder significantly outperformed the VG decoder in predicting memorability behavior (ΔPQ and 95% CI for repeated behavior: 0.31, [0.26, 0.36], H0: ΔPQ=0, p<1e-4, for novel behavior: 0.32, [0.28, 0.36], H0: ΔPQ=0, p<1e-4.) **d)** Histogram showing the angular location on the MB-VG plane where, in each cross-validation split, the maximum prediction quality was reached (across 1000 cross-validation splits), compared to a shuffled null model where, in each cross-validation split, memorability labels of each image were randomly reassigned. **e)** Total familiarity performance when decoding at axes swept along the MB-VG plane.

**Figure S7:**
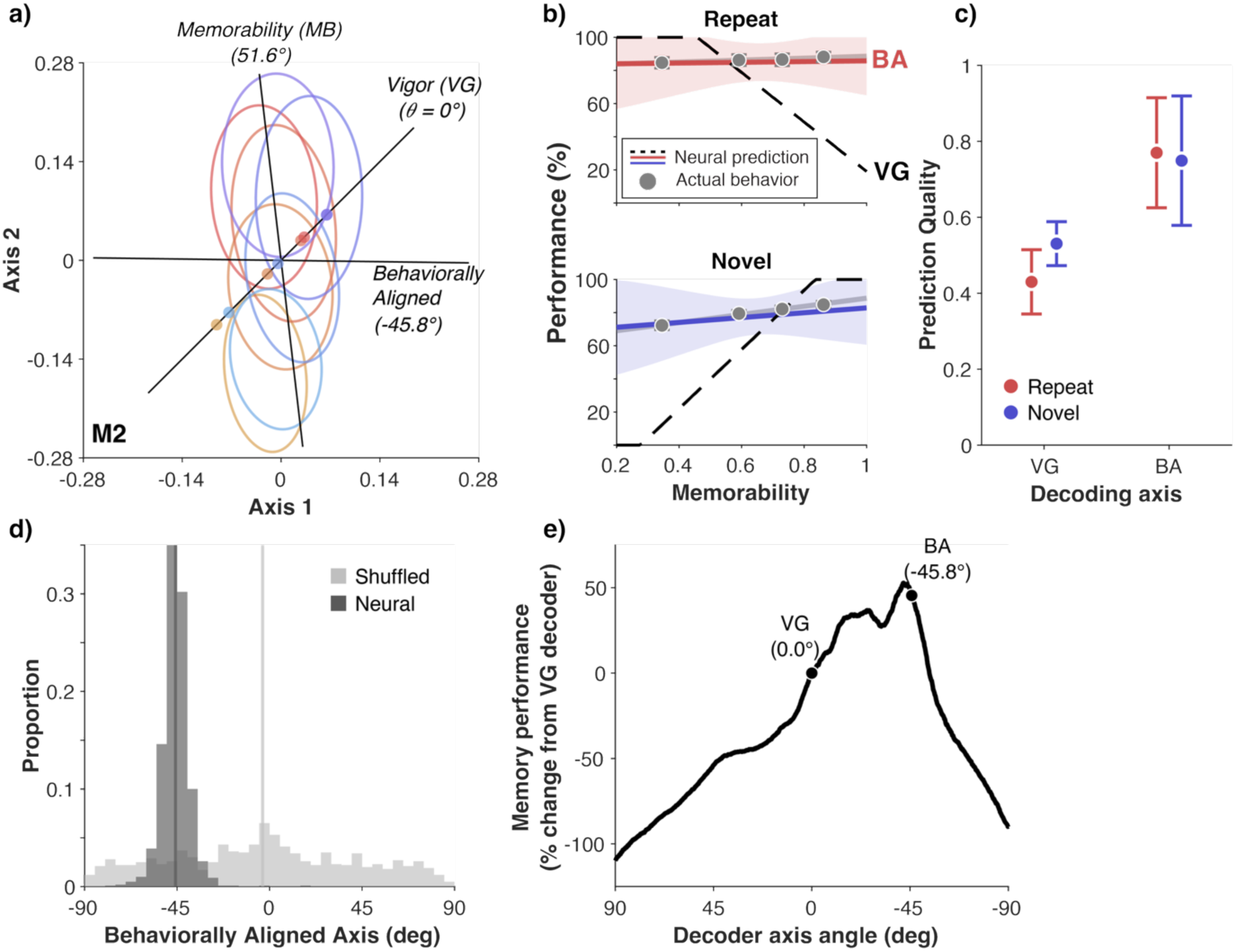
Memorability behavior is linearly decodable from ITC (M2). Same conventions as Figs 3 (top row) and S9 (bottom row). **a)** Projections of ITC neural data onto a plane defined by an axis corresponding to vigor decoder (where every unit is weighted equally, ‘Vigor (VG)’) and a linear memorability decoding axis optimized to decode high versus low memorability images, ‘Memorability (MB)’. **b)** Comparison between observed behavior (gray) and neural predictions for the vigor decoder axis (dashed line) and the behaviorally aligned decoding axis (BA, red and blue lines correspond to repeated and novel behavior, respectively). **c)** Comparison between prediction quality when decoding behavior using the vigor (VG) axis versus the behaviorally aligned (BA) axis using the same fully held-out data as described above. Decoding along the VG axis produced poor predictions of memorability behavior (repeated PQ mean +/− se: 0.43 +/− 0.08, novel PQ: 0.53 +/− 0.06). Decoding along the BA axis produced good predictions of both repeated and novel behavior when tested on fully held-out data that was not used to train either the memorability decoder or to find the BA axis (repeated PQ mean +/− se: 0.75 +/− 0.17, novel: 0.77 +/− 0.14). A bootstrap analysis (see Methods) revealed that the BA decoder significantly outperformed the VG decoder in predicting memorability behavior (ΔPQ and 95% CI for repeated behavior: 0.34, [0.29, 0.39], H0: ΔPQ=0, p<1e-4, for novel behavior: 0.20, [0.15, 0.24], H0: ΔPQ=0, p<1e-4). **d)** Histogram showing the angular location on the MB-VG plane where, in each cross-validation split, the maximum prediction quality was reached (across 1000 cross-validation splits), compared to a shuffled null model where, in each cross-validation split, memorability labels of each image were randomly reassigned. **e)** Total familiarity performance when decoding at axes swept along the MB-VG plane.

**Figure S8:**
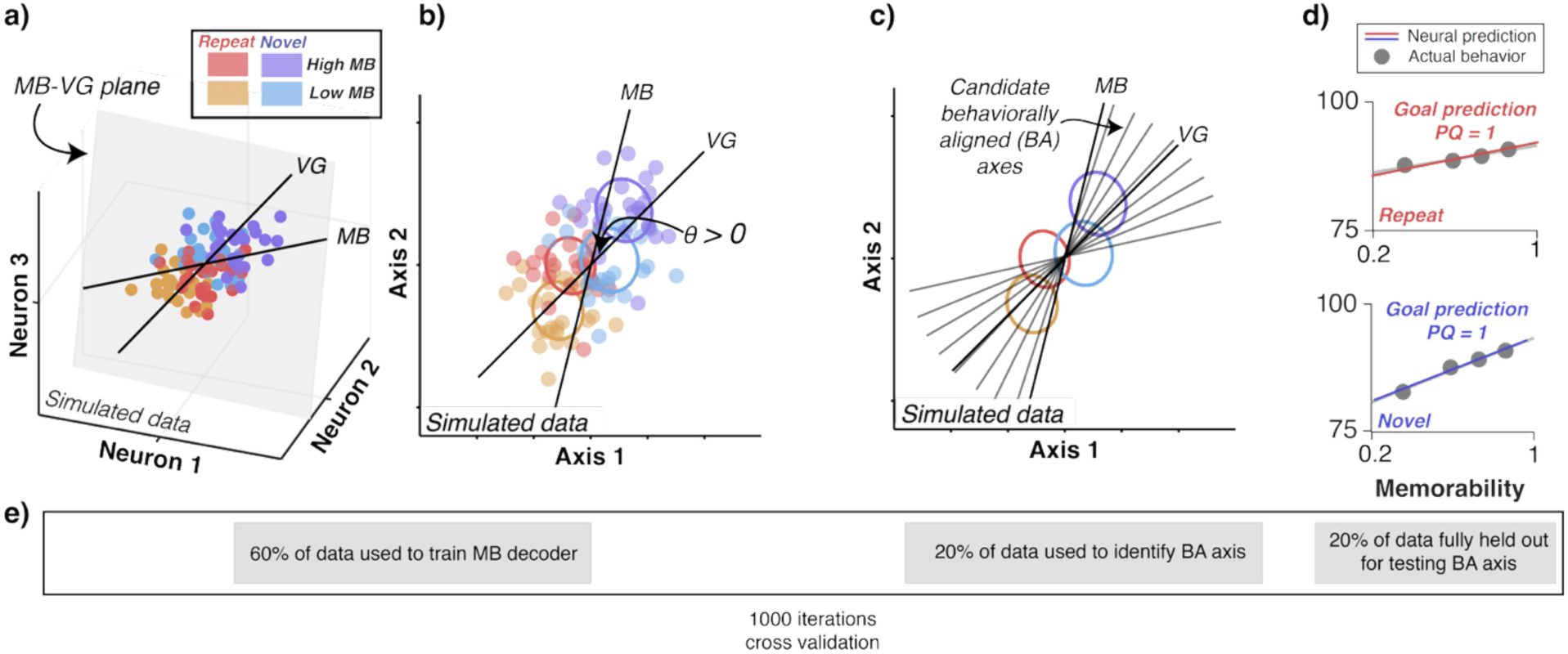
Visualization of how the behaviorally aligned decoder axis is found. **a)** Plot of the coordinate locations in the neural state-space for simulated responses from 3 synthetic neurons to 100 simulated “images” varying in memorability. For this simulation, other than memorability and familiarity, the expected firing rate of all neurons was the same to all images, and some noise was added (approximately 1/3 the amount expected by a Poisson process, to make visualization easier). The two decoders defined here in the 3-neuron space are defined by the same rules as in Fig. 3: the vigor (VG) decoder axis weights each neuron equally and positively, and the memorability (MB) decoder is trained to separate between high- and low-memorability images. The plane formed by the two non-orthogonal decoder axes is shown in this space in gray. While the example here is a 3-dimensional space corresponding to three example neurons, in a space of any dimensionality, a linear decoder axis is a 1-dimensional line, and two different lines always form a 2-dimensional plane. **b)** Projection of synthetic data onto the plane formed by the decoder axes. **c)** Ovals overlaid on the spike counts represent 2-D histograms on the MB-VG plane with axes defined by one standard deviation (as in Fig. 3a). This sets up a geometry that can be tested for whether it can align with the specific observed memorability behavior by decoding along other axes in this plane, with axes on the MB-VG plane being weighted linear combinations of MB and VG. **d)** The goal of this process is to find which, if any, decoder axis (gray lines in **c**) predicts behavior aligned with observed behavior – a prediction quality (PQ) of 1. This axis can then be interrogated with the fully held out data for whether it can predict memorability behavior. To generate predictions of behavior and determine the final PQ shown in Fig. 3, data held out from finding the MB-VG plane and the BA axis was projected onto the identified decoding axis for each cross-validation iteration and the PQ and predicted behavior were then averaged to get the result reported in Fig. 3. **e)** Schematic of cross-validation scheme. Gray rectangles arranged horizontally correspond to where (in the above plots) different subsets of each cross-validation iteration is used. The values reported in Fig. 3 are averages across 1000 cross-validation iterations.

**Figure S9:**
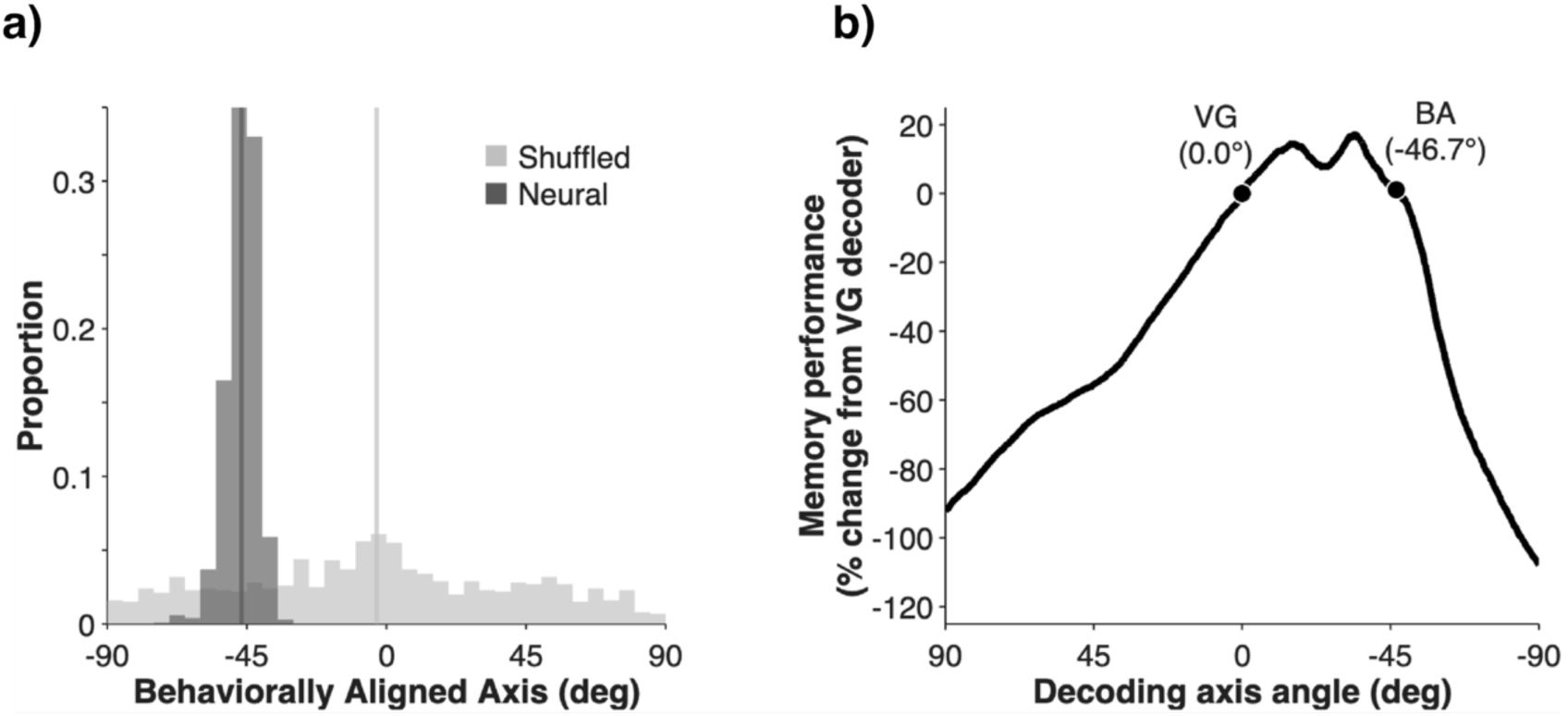
A behaviorally aligned axis is reliably found in ITC, and that axis maintains familiarity information. **a)** Histogram showing the angular location on the MB-VG plane where, in each cross-validation split, the maximum prediction quality was reached (across 1000 cross-validation splits), compared to a shuffled null model where, in each cross-validation split, memorability labels of each image were randomly reassigned. **b)** Total familiarity performance when decoding at axes swept along the MB-VG plane, compared to the VG decoding axis.

**Figure S10:**
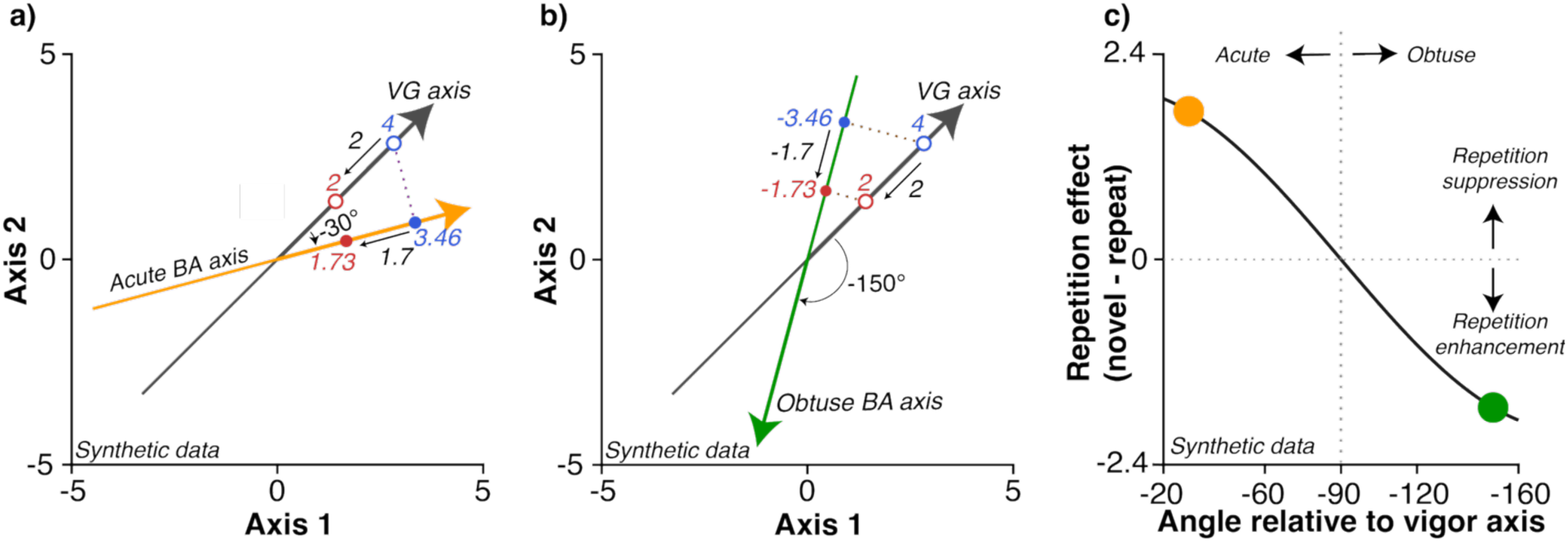
Visualization of why projections onto an acute angle retain repetition suppression, but obtuse angles invert it. **a)** Here, the firing evoked by a single, synthetic “image” is shown along the vigor (VG) axis with the novel firing plotted in blue and repeated firing in red. Repetition effect is computed by the firing rate for (novel – repeated) and a positive effect means repetition suppresses firing, while a negative effect means repetition enhanced firing. Along the VG axis, repetition effect is 4 – 2 for a repetition effect of 2 in the direction of suppression. Projected along an acute (30 deg) candidate behaviorally aligned (BA) axis, the repetition effect is 3.46-1.73 for a repetition effect of 1.7 in the direction of suppression. **b)** Projected along a candidate behaviorally aligned axis that is obtuse to the VG axis, at 150 deg, the repetition effect is −3.46 – (−1.73) for a repetition effect of −1.7 with the negative sign indicating enhancement (where repeat firing is greater than novel firing). Note that the readout takes place with respect to the direction of the axis, not the coordinate plane. **c)** Repetition effect as a function of angle relative to the VG axis. Values switch from positive (repetition suppression) to negative (repetition enhancement) at the axis orthogonal to the VG axis. The points marked on the figure in orange and green correspond to the acute axis plotted in panel a and the obtuse axis in panel b, respectively.

**Figure S11:**
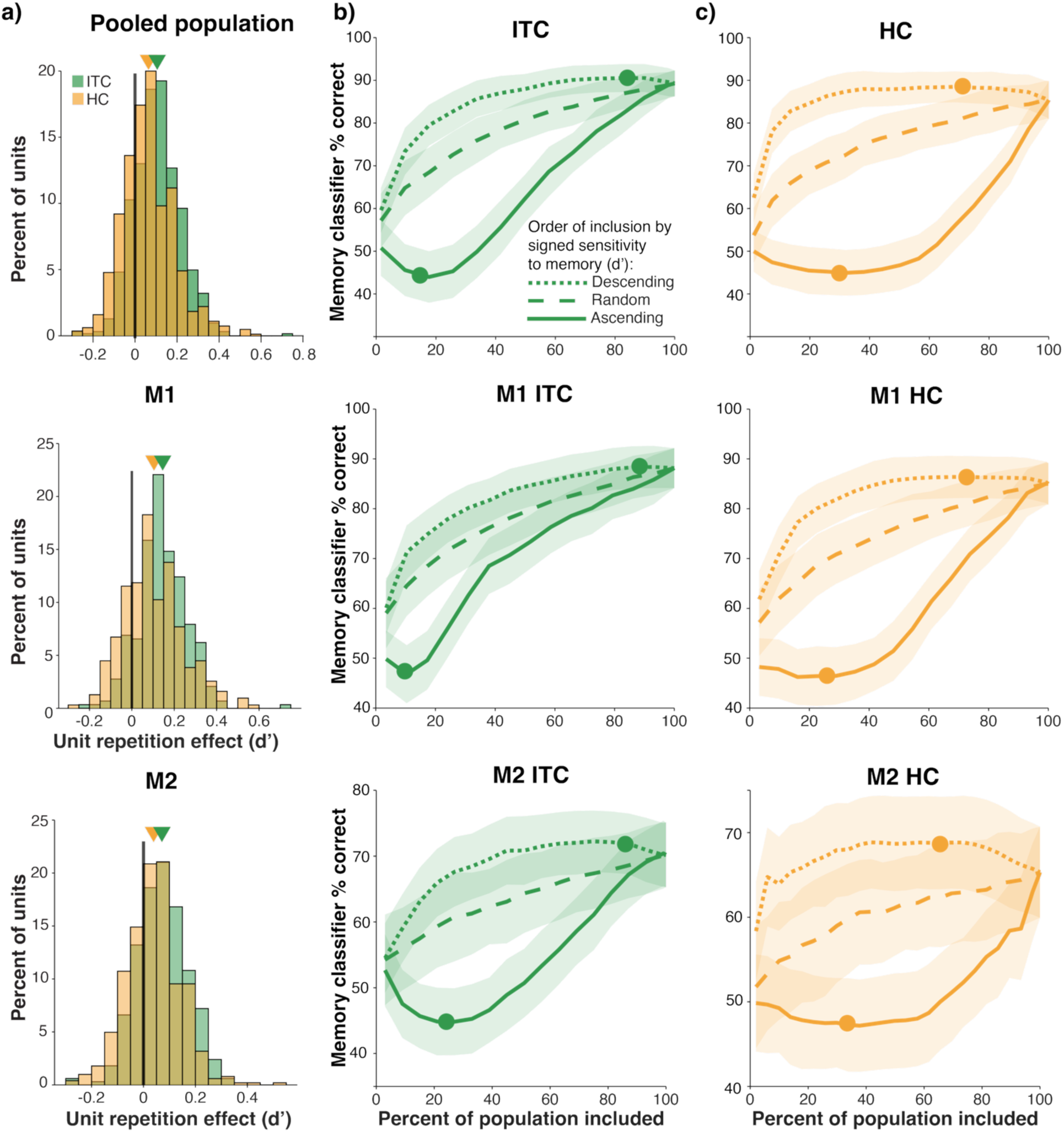
Familiarity is reflected as repetition suppression in both ITC and HC. **a)** Histograms of unitwise repetition effect (where a positive number means a unit was suppressed by familiarity and a negative number means it was enhanced). Top row shows the pooled data and the second and third rows show data analyzed for each monkey individually. Statistical tests for the top row are in the body of the paper. For M1 ITC mean = 0.14, s.d. = 0.116, t(289) = 21.2, p = 1.1e-60. For M1 HC mean = 0.10, s.d. = .141, t(311) = 12.9 p = 7.7e-31. For M2 ITC mean = 0.071, s.d. = 0.116, t(332) = 13.5, p = 1.8e-33. For M2 HC mean = 0.039, s.d. = 0.104, t(502) = 8.36, p = 6.2e-16. **b-c)** To determine if the small fraction of repetition-enhanced units observed in a) were contributing to familiarity, we performed a ranked d’ analysis. Here, we trained a weighted linear decoder applied to neural data in each population (an FLD decoder, see Methods). Note that unlike the ‘spike count decoder’ sketched in Fig. 1, this decoder is capable of negatively or positively weighting each unit to account for enhancement or suppression based on how each unit is affected by repetition. Units were ranked by their signed d’ of familiarity (i.e., in the “descending” order of inclusion, units with the most repetition suppression received the highest ranks, with repetition enhanced units with negative d’s receiving the lowest ranks) and cumulatively included portions of the population, testing for familiarity performance of each subpopulation. Broadly, the pattern between ITC and HC is similar and supports a hypothesis where familiarity information is represented in the suppressed units in both brain areas. When including units starting with the lowest d’ first (i.e., the repetition enhanced units; solid line), familiarity decoder performance in both brain areas did not exceed chance (50%) until after repetition suppressed units were included in the population. Similarly, in both populations, a large amount of the decoder performance could be accounted for with a small number of the highly repetition suppressed units (dotted lines). Unit ranks were determined from their responses to the training set of images for each cross-validation split. As such, the below chance performance from the most repetition enhanced units (ascending, solid line) suggests that these units were not truly enhanced by repetition, but more likely to have appeared so due to noise (i.e., based on the training sets in each iteration, the units appeared repetition enhanced, but when tested against the images in the test set, they were actually, on average suppressed, reflected in decoder performance dipping slightly below chance level). For this decoder, the memorability alignment of the pseudopopulation was shuffled for both training and testing to minimize the effects of the ‘paradox of memorability’, raising raw performance of the decoder compared to the unshuffled versions used in the memorability adjusted linear scheme (see Methods). Error shadows depict the standard deviation of decoder performance across 200 cross-validations at each population size. Analysis window for each brain area was 300ms-500ms following image presentation. Filled circles in the ranked analyses indicate where d’ = 0, marking the transition point in the ranked analyses between repetition suppression and repetition enhancement. The proportion of the population with a d’ below 0 was 18% for ITC and 31% for HC in the pooled population (M1: 11% for ITC, 25% for HC; M2: 23% for ITC, 35% for HC).

**Figure S12:**
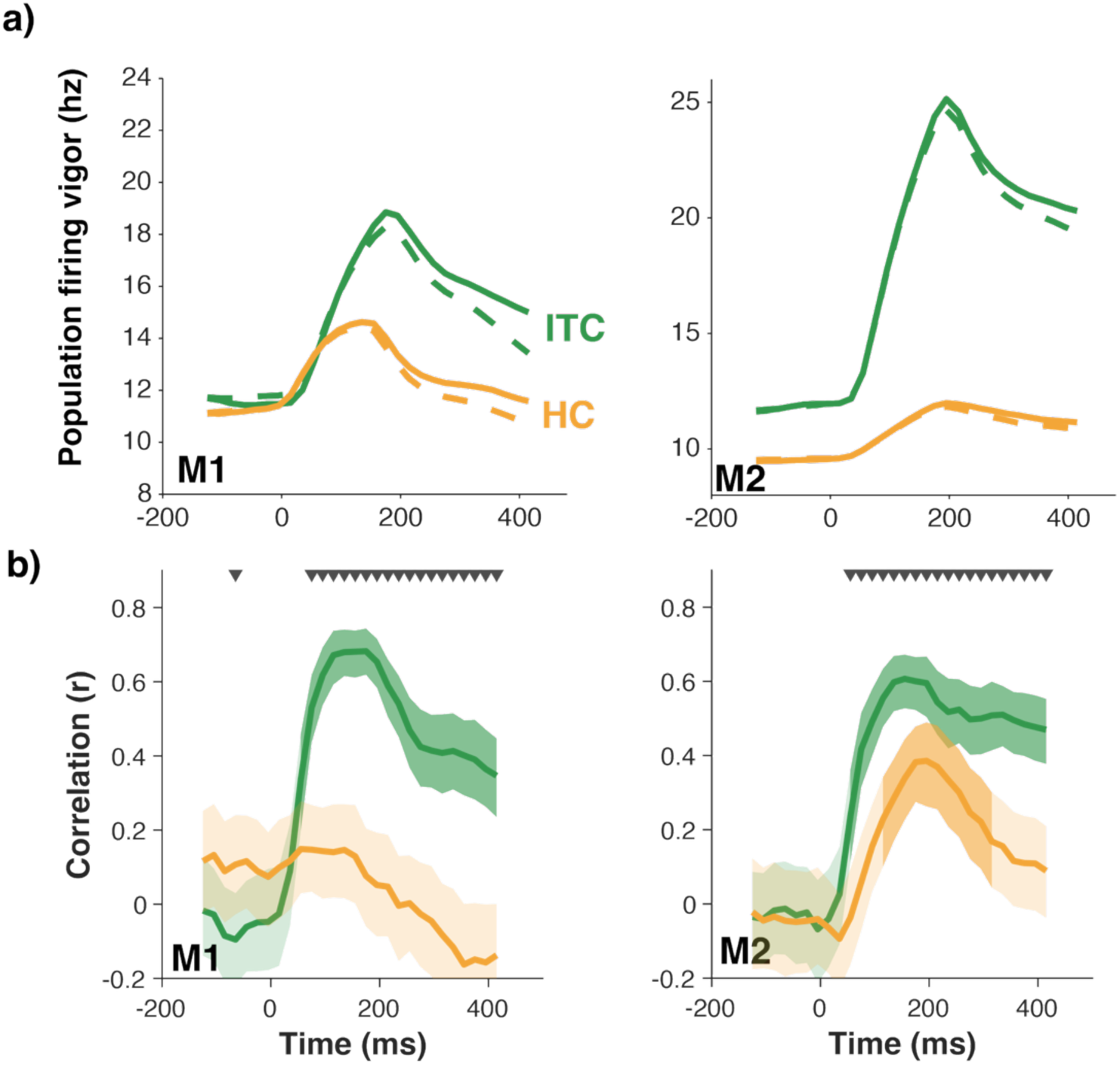
Temporal dynamics of firing and memorability in ITC and HC in individual monkeys. Same conventions as Fig. 4a and Fig. 4c. **a)** Peri-stimulus plot of firing across all trials. Solid lines indicate firing for novel images and the dashed line for repeated. **b)** Pearson’s correlation between spiking vigor across all units for repeated images with image memorability, plotted as a function of time from image onset. Shaded regions reflect 95% confidence intervals, computed by bootstrapping with 1000 resamples. Darker shadows indicate times when the correlation is significantly different from zero (p < 0.05, corrected for false discovery across time windows). Black triangles indicate timepoints when the correlations in ITC and HC were significantly different (bootstrapping, H0: Δr = 0; p<0.05 corrected for false discovery across time windows).

**Figure S13:**
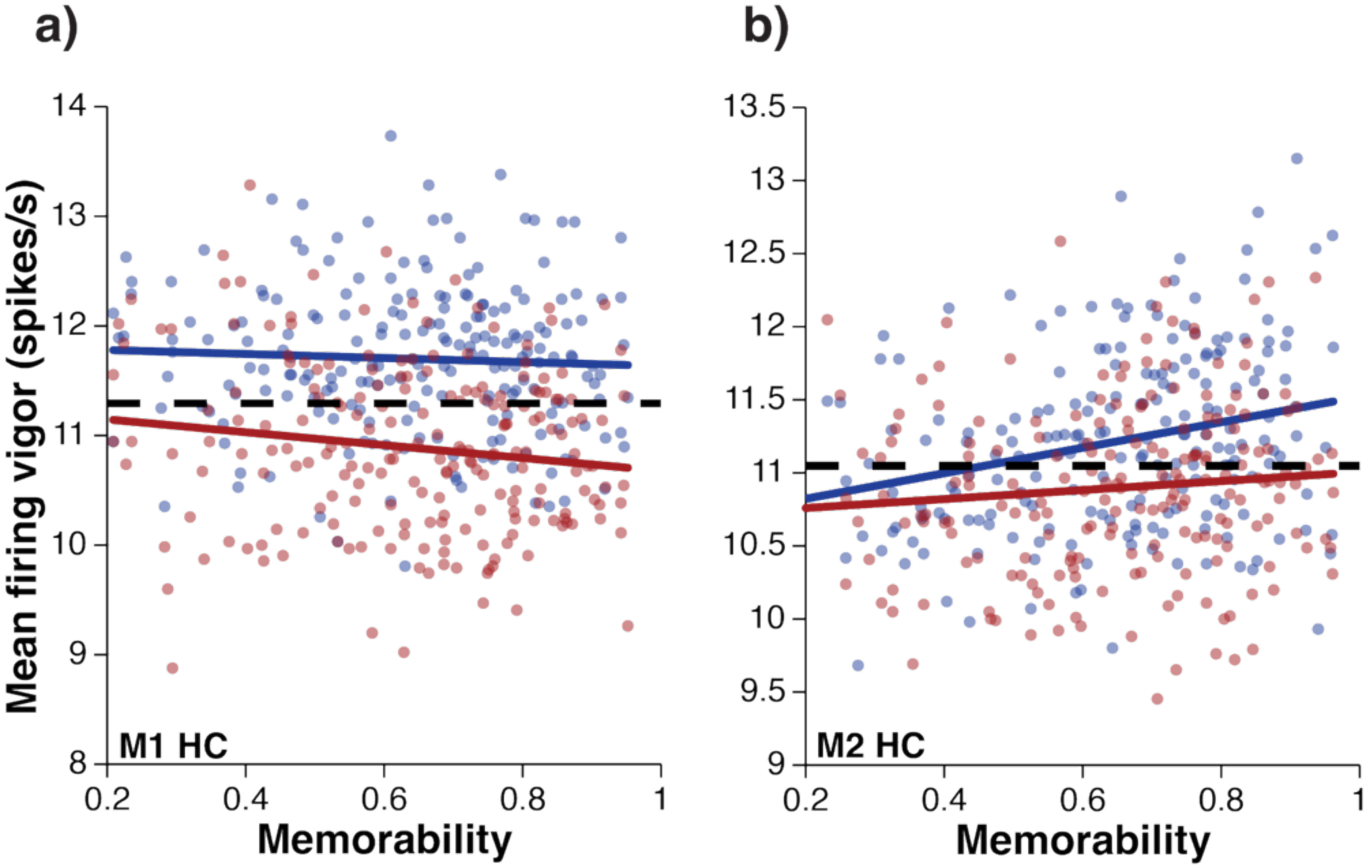
**a)** For monkey 1’s HC, the correlation between memorability and vigor for novel images was r(206) = - 0.05, p = 0.48 and for repeated images r(206) = −0.14, p = 0.04. **b)** For monkey 2’s HC, the correlation between memorability and vigor for novel images was r(206) = 0.28, p = 3.43e-5 and for repeated images r(206) = 0.11, p = 0.13. Data were analyzed in the 300-500ms window following stimulus presentation. Testing for attenuation of MB between ITC and HC in each monkey: For monkey 1 novel ITC β=5.234, HC β=-0.183 (H0: Δβ = 0 p < 1e-5), for repeated ITC β=4.254, HC β=-0.588 (H0: Δβ = 0 p = 2e-4). For monkey 2, novel ITC β=6.710, HC β=0.874 (H0: Δβ = 0 p = 0.023), for repeated ITC β=5.916, HC β=0.308 (H0: Δβ = 0, p = 0.021).

**Figure S14:**
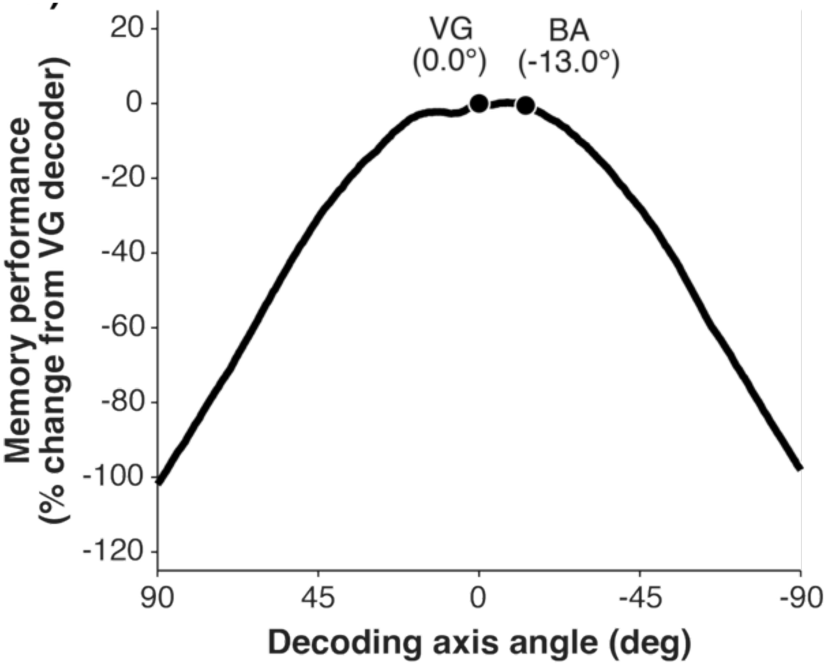
The behaviorally aligned axis in HC maintains familiarity information compared to the vigor decoder. Total familiarity performance when decoding at axes swept along the MB-VG plane. Plotted with the same conventions as Fig. S9b.

**Figure S15:**
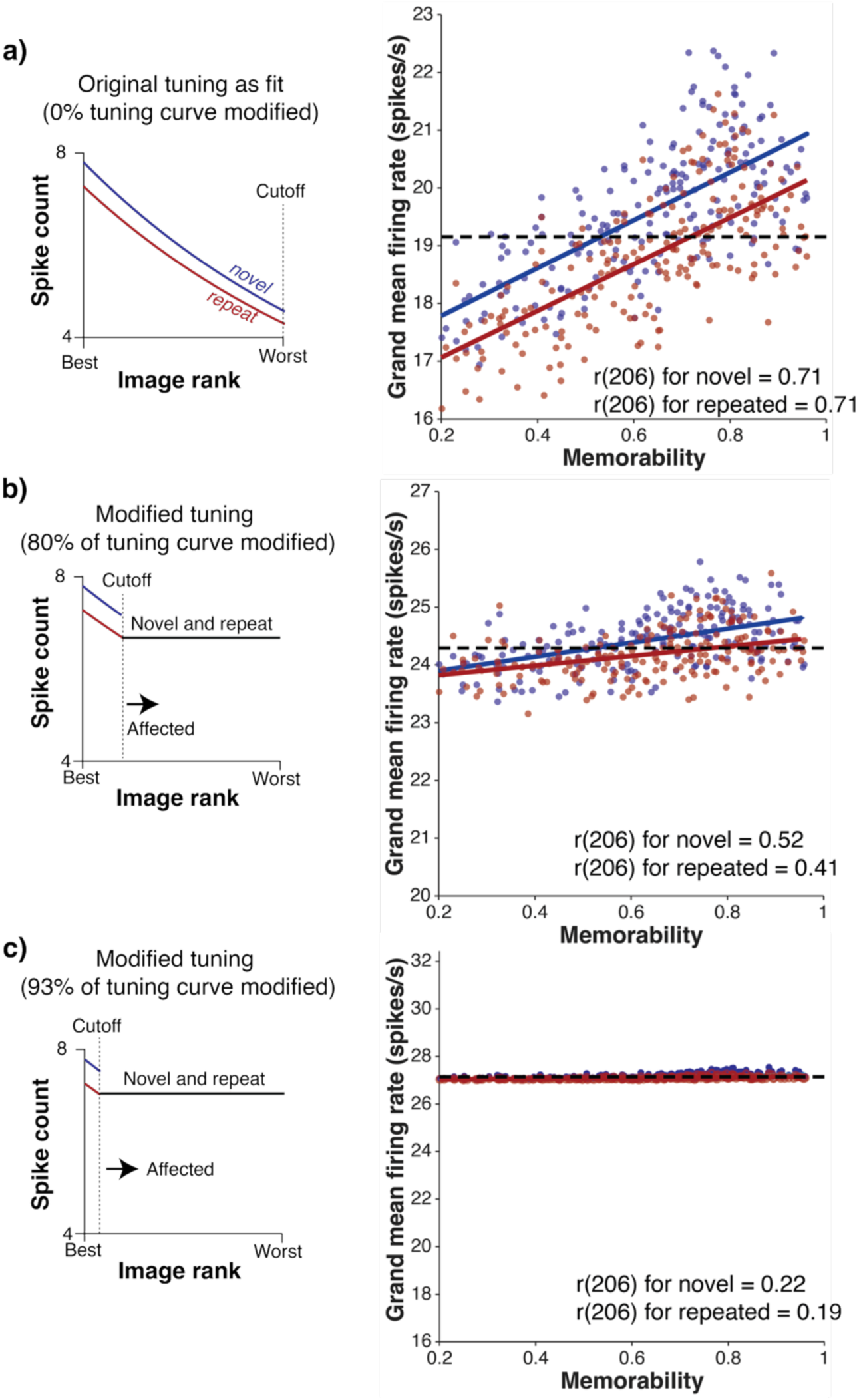
Example synthetic ITC populations. **a)** Example fit tuning curve (left) used to generate synthetic population. Equivalent analysis to Fig. 2a on synthesized data (right). The synthetic population was generated by fitting exponential tuning curves to each unit and using a Poisson process to simulate spikes (see Methods for details). **b-c)** Populations generated based on modified tuning curves at 80% tuning thresholded (b) and 93% of tuning thresholded (c).

